# Structural dynamics of the β-coronavirus M^pro^ protease ligand binding sites

**DOI:** 10.1101/2021.03.31.437918

**Authors:** Eunice Cho, Margarida Rosa, Ruhi Anjum, Saman Mehmood, Mariya Soban, Moniza Mujtaba, Khair Bux, Sarath Dantu, Alessandro Pandini, Junqi Yin, Heng Ma, Arvind Ramanathan, Barira Islam, Antonia S J S Mey, Debsindhu Bhowmik, Shozeb Haider

## Abstract

β-coronaviruses alone have been responsible for three major global outbreaks in the 21^st^ century. The current crisis has led to an urgent requirement to develop therapeutics. Even though a number of vaccines are available, alternative strategies targeting essential viral components are required as a back-up against the emergence of lethal viral variants. One such target is the main protease (M^pro^) that plays an indispensible role in viral replication. The availability of over 270 M^pro^ X-ray structures in complex with inhibitors provides unique insights into ligand-protein interactions. Herein, we provide a comprehensive comparison of all non-redundant ligand-binding sites available for SARS-CoV2, SARS-CoV and MERS-CoV M^pro^. Extensive adaptive sampling has been used to explore conformational dynamics employing convolutional variational auto encoder-based deep learning, and investigates structural conservation of the ligand binding sites using Markov state models across β-coronavirus homologs. Our results indicate that not all ligand-binding sites are dynamically conserved despite high sequence and structural conservation across β-coronavirus homologs. This highlights the complexity in targeting all three M^pro^ enzymes with a single pan inhibitor.

## Introduction

Coronaviruses (CoVs) belong to a family of positive-sense, single stranded RNA viruses with spherical envelope and a crown-like appearance due to their distinctive spike projections.^1, 2^ While α or β-coronaviruses infect mammals, γ and δ-coronavirus can infect birds or mammals (Figure 1A).^3^ Currently, seven CoVs have been identified that infect humans, namely human coronavirus 229E (HCoV-229E), OC43 (HCoV-OC43), NL63 (HCoV-NL63), Hong Kong University-1 (HCoV-HKU1), severe acute respiratory syndrome coronavirus (SARS-CoV), Middle East respiratory syndrome coronavirus (MERS-CoV) and the severe acute respiratory syndrome coronavirus 2 (SARS-CoV2).^1, 2, 4–9^ The first four are responsible for 5-30% of the common cold,^10^ while the latter three cause acute lung injury, acute respiratory distress syndrome, septic shock, and multi-organ failure with a high case fatality ratio.^1, 11^ β-coronaviruses alone have been responsible for three major global outbreaks in the 21^st^ century: SARS in 2002, MERS in 2013 and COVID-19 in 2019, with a fatality rate of 10%, 34% and 3-5% respectively.^12^

**Figure 1:**
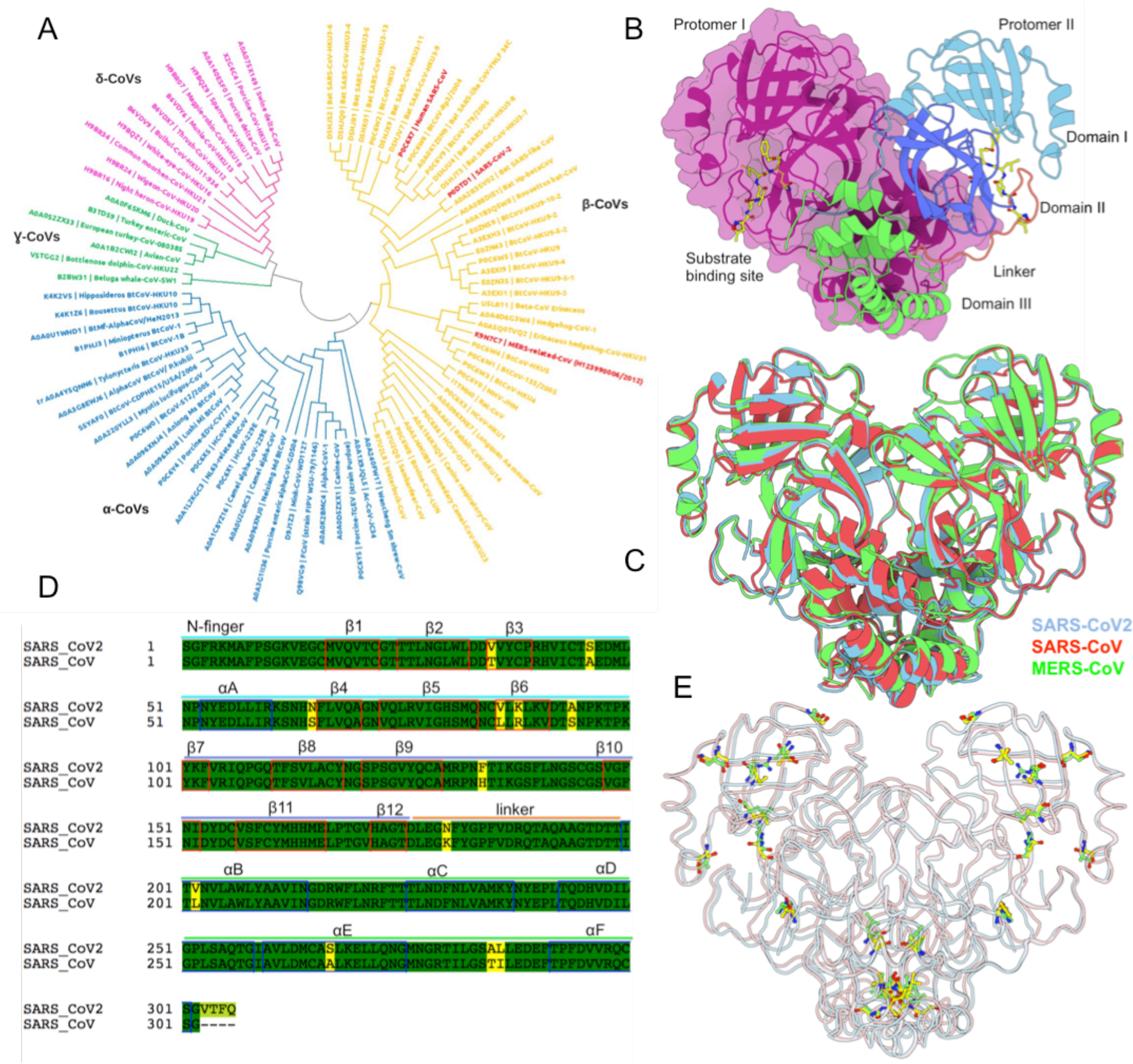
Overview of β-coronavirus M^pro^. (A) A phylogenetic tree of the α (blue), β (yellow), γ (green) and δ (pink) coronavirus family; (B) Structure of the dimeric SARS-CoV2 M^pro^ enzyme (PDB 6LU7). The two protomers are represented in two different colors; distinct structural domains in protomer II are illustrated as cartoons; (C) Comparison of SARS-CoV2 (PDB 6LU7, cyan), SARS-CoV (PDB 2C3S, red) and MERS-CoV (PDB 4YLU, green) crystal structures; (D) Sequence alignment between SARS-CoV2 and SARS-CoV highlighting the position of 12 dissimilar residues in yellow. The structural elements have been annotated on the sequence and (E) Spatial position of the dissimilar residues (yellow, SARS-CoV2; green, SARS-CoV) highlighted on the M^pro^ structure.

Coronavirus have the largest genome amongst any RNA viruses, with a size ranging between 26-32 kb.^13, 14^ The replication cycle of CoVs is initiated by the spike protein attaching to the host receptor, inducing fusion events that allow viral entry into the host cell.^15^ Once released inside, the viral genome is expressed into a series of proteins using multiple open reading frames (ORFs). In the SARS-CoV2 genome, 23 unannotated viral ORFs have been identified and include upstream ORFs that are likely to have a regulatory role; several in-frame internal ORFs within existing ORFs, resulting in N-terminally truncated products, as well as internal out-of-frame ORFs, which generate novel polypeptides.^16^ Of these, two overlapping ORFs (ORF1a and ORF1b), which makes up of 2/3^rd^ of its genome, are translated into two large polyproteins (pp1a and pp1ab). The remaining genome is transcribed into conserved structural (spike, envelope, membrane and nucleocapsid) and accessory proteins that are not essential for virus replication but have a role in pathogenesis.^17^

The pp1a and pp1ab are processed by two conserved viral proteases, 3-chymotrypsin-like cysteine protease (3CL^pro^ or M^pro^) and papain-like protease (PL^pro^), into 16 non-structural proteins (Nsp1-16), which are essential for viral replication and transcription.^18^ M^pro^ is encoded by Nsp5 and auto-cleaved from polyproteins to produce a mature enzyme. The M^pro^ enzyme then cleaves 11 downstream non-structural proteins important for viral replication, thereby making M^pro^ an essential protein for the viral life cycle.^19^ The substrate recognition sequence of M^pro^ at most sites is x-(L/F/V)Q↓(G/A/S)-x (x = any amino acid; ↓ cleavage site), where the glutamine prior to cleavage site is essential.^20^ No human protease with similar cleavage specificity is known. Thus compounds that target this cleavage site on M^pro^ will have little or no impact on human cellular proteases.^21^ This makes M^pro^ an attractive drug target.

The SARS-CoV2 M^pro^ structure is a homodimer, with each protomer (residues 1-306) composed of three domains (Figure 1B). Domain I (residues 8-101) consists of 6 β-strands (β 1-6) and one α-helix (α-helix A), while domain II (residues 102-184) consists of 6 β-strands (β 7-12). The β-strands form an antiparallel β-barrel structure in each domain and uses a long linker loop (residues 185-200) to connect to domain III (residues 201-303), which has five α-helices (α-helix B-F) arranged in a compact antiparallel globular cluster.^22^ The substrate-binding site is present in a cleft between domains I and II and buries the C145-H41 catalytic dyad. During the hydrolysis reaction, C145 acts as a nucleophile, while H41 acts as a base catalyst. An oxyanion hole formed by the backbone amido groups of G143 and C145 stabilizes the partial negative charge developed at the substrate cleavage bond.^23^ The substrate-binding site consists of four subsites (S1’, S1, S2, S4) that define enzyme specificity for glutamine in the substrate.^22^ Moreover, in the homodimer structure of the M^pro^ enzyme, the N-finger (residues 1-7) of one protomer is squeezed between domain I and II to shape the substrate-specificity pocket. This shows the importance of dimerization and N-finger orientation for substrate specificity and catalysis.^24^ Further structural analysis of the SARS-CoV2 M^pro^ identified that domain I and II are connected via seven residues (D92-P99) that contribute to the substrate-binding site.^21, 22^ Domain III contributes to the proteolytic activity via dimerization of the M^pro^ enzyme.^22^ Dimerization is important because monomeric M^pro^ does not exhibit any catalytic activity.^25^ Since M^pro^ is a symmetric homodimer, two copies of the ligand binding sites are present, one on each protomer.

A comparison of the SARS-CoV2, SARS-CoV and MERS-CoV M^pro^ sequences revealed that SARS-CoV2 is 96% similar to SARS-CoV and 51% with MERS-CoV (Figure S1). A structural superimposition of the all M^pro^ enzymes displayed an overall RMSD of 0.85 Å (± 0.16 Å), with a very high degree of structural conservation around the catalytic dyad in the substrate-binding site, suggesting very similar substrate recognition profiles amongst these proteins (Figure 1C). The difference between SARS-CoV2 and SARS-CoV M^pro^ is 12 amino acids in each protomer (Figure 1D/E). These residues have been illustrated in Figure S2.

The past year has seen a dramatic progress in SARS-CoV2 research (covid19primer.com). Significant efforts have gone into the design of M^pro^ inhibitors that target the substrate binding pocket.^21, 22, 26, 27^ This also includes inhibitor design via various *in silico* methods.^28–33^ Recent progress on M^pro^ inhibitors have been reviewed elsewhere.^34–36^ Of considerable note is the COVID Moonshot project, which generates data via open science discovery of M^pro^ inhibitors by combining crowdsourcing, high-throughput experiments, computational simulations and machine learning.^37^ This alone has generated over 258 structures of fragment and lead-like molecules in complex with the M^pro^ protease (www.covid.postera.ai/covid). Other large-scale efforts using crystallographic screening of fragments and drug repurposing libraries have identified allosteric drug binding sites.^38, 39^

Over 270 crystal structures of SARS-CoV2 M^pro^ are present in the protein data bank (PDB), including apo and co-complexes with inhibitors (Table S1). Additional similar data is also available on SARS-CoV (Table S2) and MERS-CoV (Table S3) M^pro^ enzymes. However, to date, no comprehensive, consolidated, comparison of ligand binding sites and their complexes determined using X-ray crystallography has been reported. In this study, we map non-redundant ligand binding sites from all crystal structures of SARS-CoV2 M^pro^ available in the PDB. We then carry out 25 µs of adaptive molecular dynamics (MD) simulations on the apo M^pro^ structure of SARS-CoV2, SARS-COV and MERS-CoV and explore the conformational dynamics of the M^pro^ enzymes using a deep learning approach, namely a variational auto encoder with convolutional filters (CVAE),^40^ and investigate the structural conservation of the ligand binding sites using Markov state models (MSM). Further, we annotate each binding site with a measure of correlated evolution at the residue level. Our results highlight that even though with a structural overlap of <1 Å, the conformational dynamics of SARS-CoV2, MERS-CoV and SARS-CoV are very different. A persistence analysis and comparison of the structural conservation of the ligand binding sites in β-coronavirus homologs highlight the complexity in targeting all three M^pro^ enzymes with a single pan inhibitor.

## Results

### Mapping the binding sites

The PDB was searched for β-coronavirus M^pro^ entries. A total of 271 SARS-CoV2 structures were identified. Out of these, there are 38 structures with no ligands and were excluded from any further study. The remaining 233 structures were downloaded for detailed structural analysis and are listed in Table S1-S6. The key interacting residues between the inhibitors and M^pro^ were mapped (Figure S3). In total, 22 different binding sites were identified. These have been labelled A-V in Figure 2 and listed in Table 1. A detailed structural description of the binding site is provided in the supporting information.

**Table 1.** SARS-CoV2 M^pro^ ligand binding sites.

| Binding site | Binding site residues | Ligand ID | PDB (representative structure) | Number of ligands |
| --- | --- | --- | --- | --- |
| A | T25, H41, M49, Q189, H164, A191, C145, N142, S144, H163, E166, H172, P168 | N3 | 6LU7 | 185 |
| B | I78, G79, H80, S81, K88, K90 | K1Y | 5RFC | 5 |
| C | H80, I59, E55, L58, S81 | O0S | 5RE6 | 2 |
| D | V186, R188, T190, A191, Q912 | SFY | 5RF8 | 1 |
| E | F103, E178, D176, R105, V104 | HV2 | 5RF5 | 4 |
| F | R105, Y182, G183, P184, F134, P108, Q107, I106 | LWA | 5REG | 1 |
| G | H172, G138, R4, G2, F3 L282, K137, G170, V171 | T5D | 5RF0 | 1 |
| H | P132, T198, Y196, E240, Y239, M235, N238 | S7V | 5RGS | 4 |
| I | P241, M235, I232, N231, N228 | K1G | 5RGR | 1 |
| J | A70, V73, T93, P96, W31, K97 | T0S | 5RE7 | 1 |
| K | P96, N95, A94, D34, D33 | T6J | 5RFD | 3 |
| L | P96, T98, P99, K100, D155, K12, K97 | S7D | 5RF9 | 3 |
| M | Y118, L141, S123, F8, Q127, D295, A7, D155, R298, Q299, M6 | JGY | 5RFA | 2 |
| N | Q110, F295, V297, T292, P293, I249, P252 | 6SU | 5REF | 2 |
| O | N27, G278, R279 | JGP | 5REA | 1 |
| P | L287, A285, M276, G275, N274, L271, Q273, Y237, L272 | QCP | 7AXO | 2 |
| Q | N142, C300, S301, V297, P252, L253, Q256 | RMZ | 7AMJ | 5 |
| R | K100, Y101, K102, C156, D155 | UHG | 7ARF | 6 |
| S | D33, K102, Y101, K100, P99 | RVW | 7AWR | 2 |
| T | V233, Y237, K269, Q273 | X4P | 7KVL | 1 |
| U | Q83, N84, K88 | X4V | 7KVR | 1 |
| V | R4, K5, A7, V125, Y126, Q127 | XY4 | 7LFP | 1 |

**Figure 2:**
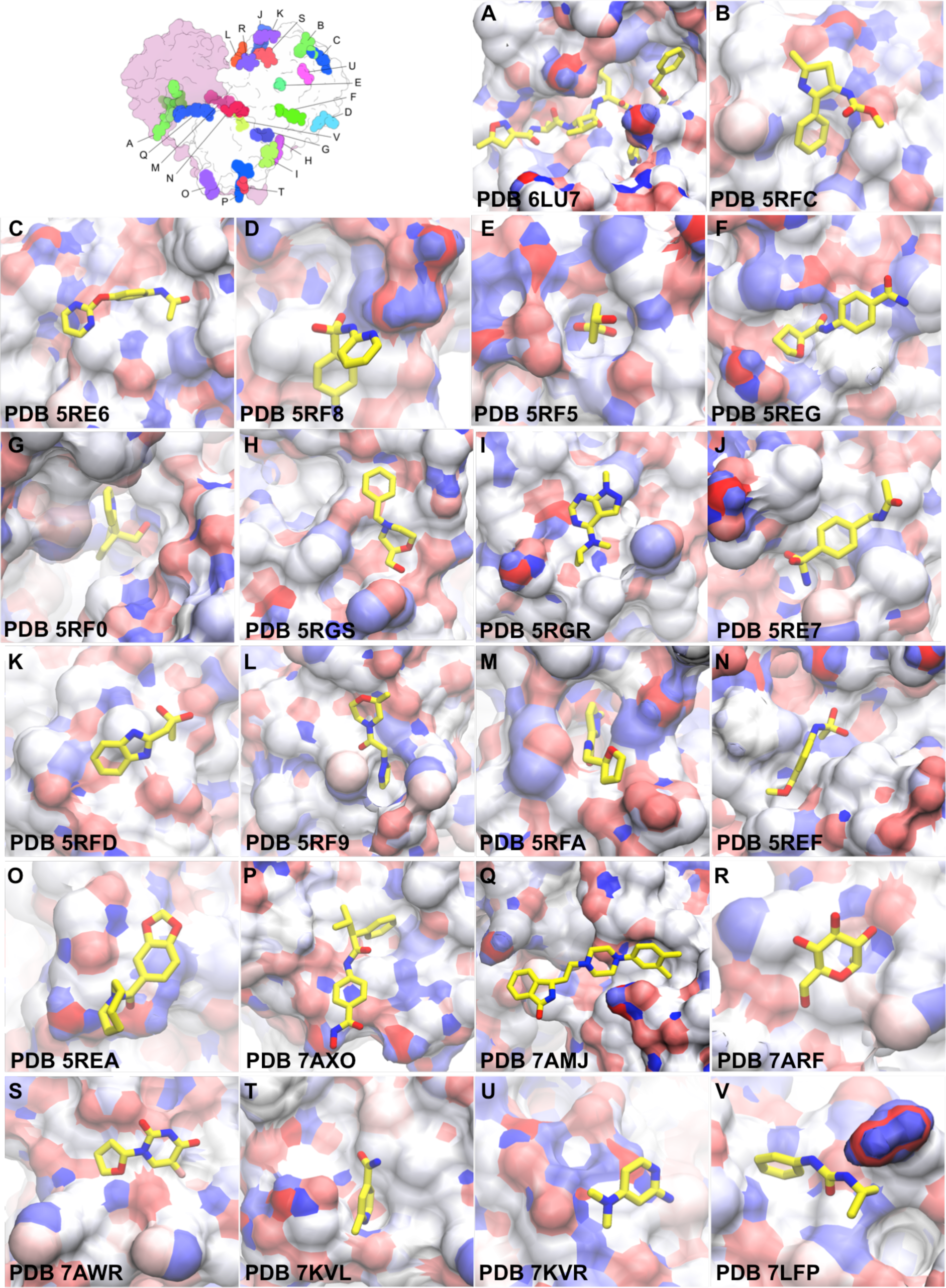
Ligand binding sites on SARS-CoV2. (A) An overview of the ligand binding sites identified from X-ray structures. While there are two copies of each binding site (one on each protomer), only one copy is illustrated. (A-V) 22 non-redundant ligand binding sites identified from various SARS-CoV2 representative structures. The surface is colored by charge.

### Structural dynamics of the β-coronavirus M^pro^ enzymes

To further understand the structural dynamics of the β-coronavirus M^pro^ enzymes, MSM-based adaptive sampling molecular dynamics (MD) simulations were conducted. These simulations have an advantage over classical MD in exploring under sampled states without a predetermined bias. The sampling and analysis mainly focused on investigating the differences in dynamics of the M^pro^ enzyme ligand binding sites. SARS-CoV2 and SARS-CoV are 96% identical, with a difference of only 24 residues out of 612. When structurally aligned, the root means squared deviation (RMSD) of the protein backbone aligned structures is 0.61 Å. This is also similar to when comparing with the MERS-CoV structure, where the sequence similarity is ∼51% and the structural alignment is 0.51 Å.

The conformational drift during the course of the simulations was assessed using Cα root mean-squared deviation (RMSD). Conventional RMSD fitting methods fail to separate regions of different stability. To resolve such regions, we used a fraction (%) of the Cα atoms for the alignment. Beyond this fraction, there is a sharp increase in the RMSD value for the remainder of the Cα atoms. At 60%, the core of the M^pro^ could be superimposed to less than 0.12 nm (1.2 Å), 0.10 nm (1 Å) and 0.09 nm (0.9 Å) for SARS-CoV, SARS-CoV2 and MERS-CoV M^pro^ structures (Figure S5A). The Cα atoms above 60% cutoff predominantly belong to the dimerisation domain III, the linker loop and the loops in domain I and II. The antiparallel β-barrel structures show the least deviation (Figure S5B-D).

### CVAE-based Deep learning analysis

CVAE can organize the conformational landscape in terms of a small number of biophysically relevant conformational coordinates from long timescale simulations.^40, 41^ In protein folding trajectories, the CVAE-inferred reduced conformational coordinates correspond to separating folded and unfolded states (and potentially transitions involving these states)^40^ and for equilibrium simulations, the CVAE provides biophysically relevant information related to conformational transitions induced by changes in hydrogen-bonds/hydrophobic interactions.^41, 42^ In this work, the CVAE was used to identify any differences in the collective conformational fluctuations within M^pro^ simulations from SARS-CoV2, SARS-CoV and MERS-CoV.

First, the CVAE model quality from the simulation trajectories was evaluated by examining the training and validation loss for various models, with latent dimension size ranging from 3 to 11 (Figure 6B). Initially, as the dimension size increases, the corresponding model compresses less and hence has more representation capability. When the latent dimension becomes too large, the model may over fit local features and introduce extra noise, and the regularizing term (Kullback-Leibler divergence) of the loss will play a bigger role. The overall loss approaches an optimal value in between those two extremes. For this dataset, the CVAE model is quite stable and robust, considering the validation loss stays close across various latent dimensions.

A latent dimension of 7 was selected based on the reconstruction loss as well as the uncertainty in the validation set. Next, t-distributed stochastic neighbor embedding (t-sne) was performed on this compressed lower-dimension data to visualize in two dimensions.^40^ Figure 3A shows the two-dimensional t-sne representation of the CVAE low-dimension data, while Figure 3B depicts three dimensions of the compression CVAE low-dimensional data. The CVAE is able to completely cluster the three different β-coronavirus M^pro^ types based on the local and global conformational dynamics (Figure 3). This is more visible in the two-dimensional representation. Here, SARS-CoV2 and SARS-CoV behave similar to each other while MERS-CoV is very different. One must note that this clustering is not evident when using traditional features (such as RMSD or native contacts) to distinguish among these three types of closely related β-coronavirus homologs, proving the sensitivity of the CVAE implementation.

**Figure 3:**
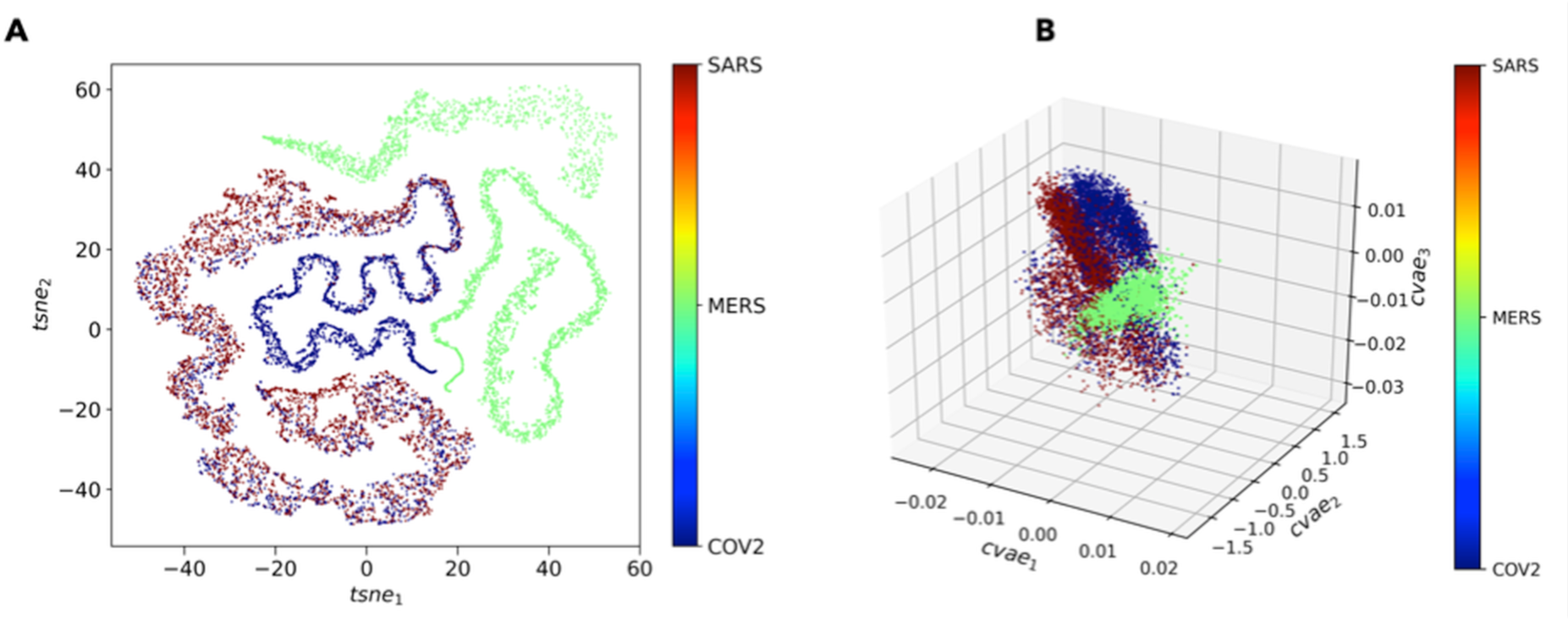
CVAE based Deep Learning Analysis. Low dimensional latent space of CVAE learnt features of the high dimensional input in (A) 2D representation and in (B) 3D representation. Original high dimensional data is transformed into distance matrix format, which is then fed to CVAE architecture. The CVAE captures the intrinsic features of the high dimensional data that are necessary to describe the original system behavior. This captured information is then shown into three-dimensional format (right) and in two-dimensional format (left) following t-sne treatment. The results show that MERS-CoV (green) dynamics is very different from SARS-CoV (red) or SARS-CoV2 (blue).

### Markov State Model

The main focus for building an MSM was to investigate how the various binding sites identified from X-ray crystallography (Table 1-3) were linked dynamically in the network of metastable states and transition probabilities among them. The choice of this method was based on the ability of MSM methods to use large ensembles of short-timescale trajectories for sampling events that occur on slow timescales.^43, 44^ The metastable states are an ensemble of structural conformations that interconvert quickly within the ensemble and slowly between them. These ensembles broadly correspond to the different basins on the free energy landscape (FEL). MSMs, provide a powerful method for detecting metastable states, calculating kinetics and free energies by integrating any number of simulations into a single statistical model.^43–48^

**Table 2.**
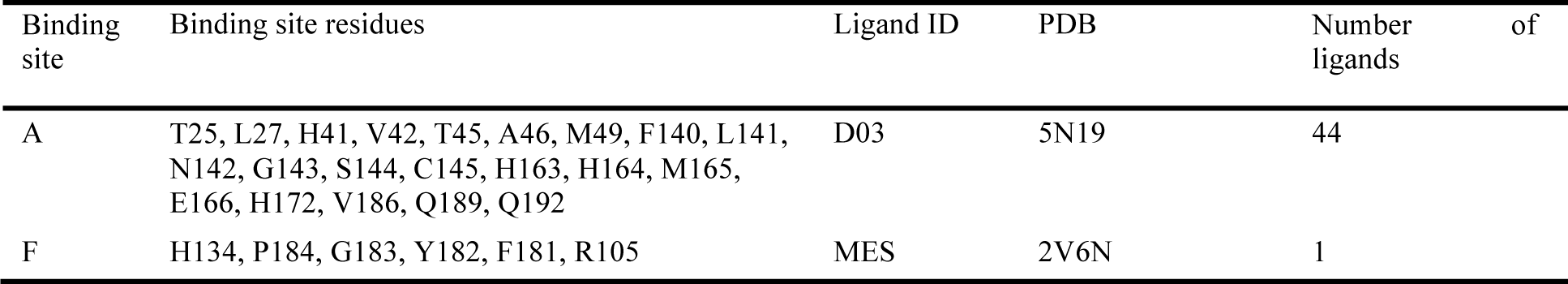
SARS-CoV M^pro^ binding sites.

| Binding site | Binding site residues | Ligand ID | PDB | Number of ligands |
| --- | --- | --- | --- | --- |
| A | T25, L27, H41, V42, T45, A46, M49, F140, L141, N142, G143, S144, C145, H163, H164, M165, E166, H172, V186, Q189, Q192 | D03 | 5N19 | 44 |
| F | H134, P184, G183, Y182, F181, R105 | MES | 2V6N | 1 |

**Table 3.**
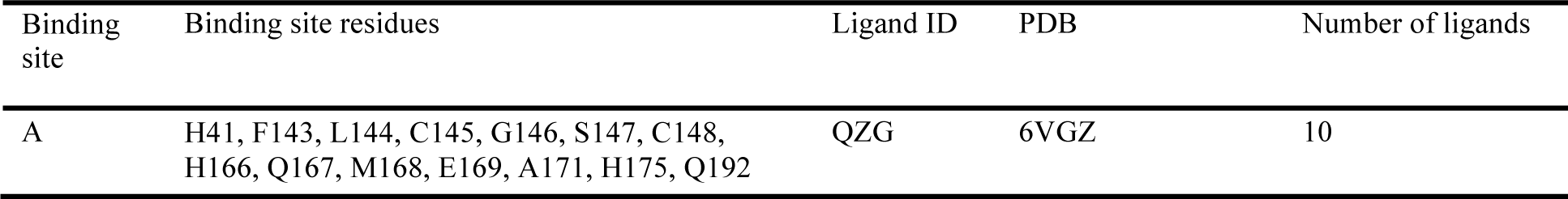
MERS-CoV M^pro^ binding sites.

We first used φ and ψ dihedral angles of the 24 residues that are dissimilar between SARS-CoV2 and SARS-CoV (Figure S2) as input data. However, this data was not sufficient to build a converged MSM. We then included φ and ψ dihedral angles of all residues and χ1 angle from the 24 residues that were different as input data to construct MSM. The dimensionality of the data was further reduced through time-independent component analysis (tICA) and the models built using PyEMMA software, from a set of 520 short 50 ns enhanced sampling MD simulations. It was possible to build converged MSM with a lag time of ≥10. Shorter lag times provide more structural detail but can underestimate the populations of important states, while simulations with longer lag times provide better population estimates but obscures intermediate states. The data was clustered into 100 microstates and their distribution on the FEL is presented in Figures S7-9. Transition pathways were then generated to identify metastable conformations. In total 5 metastable states were identified for SARS-CoV2 and 5 for SARS-CoV and 4 for MERS-CoV (Figure 4).

**Figure 4:**
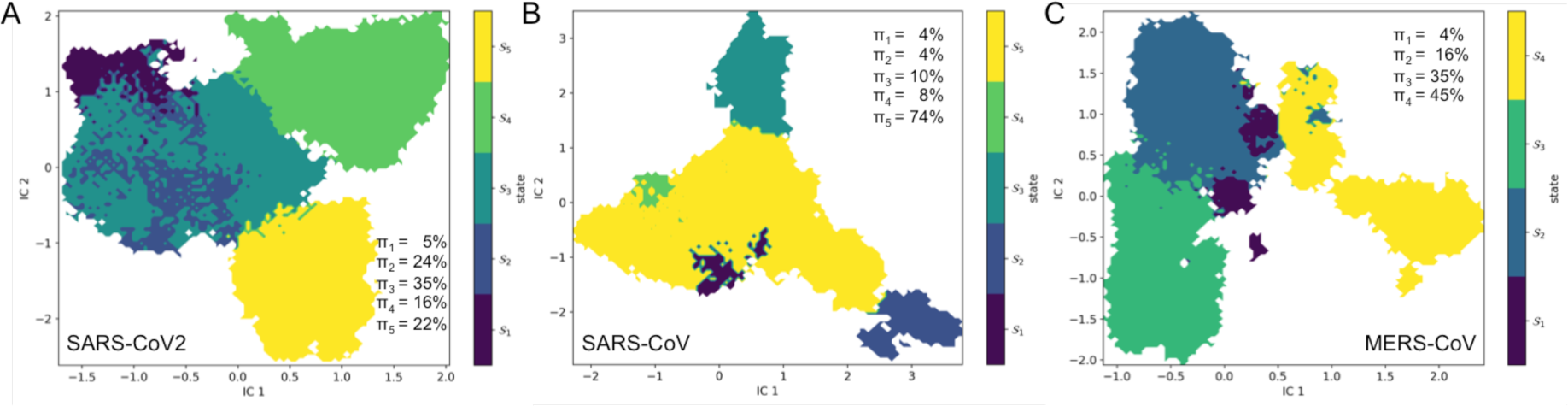
Markov State Network. Macrostate distributions of (A) SARS-CoV2 (B) SARS-CoV and (C) MERS-CoV conformations projected onto first two time-lagged independent components (ICs). The population of each state (π) is indicated in the figure. The trajectory has been aligned to state S1 and we assume this macrostate to be the crystal structure-encompassing state. The state with the highest population is classified as the dominant state. The representative metastable structures are illustrated in Figures S7-S9.

### Dynamic pocket tracking

Since many of the ligand binding sites appears together on the M^pro^ surface, we investigated the spatiotemporal evolution of the binding pockets. Protein conformations of the metastable states were searched for the presence of the experimentally reported binding sites. The site was described as open, if it could hold a minimum of five water molecules, which was a coarse equivalent of a small fragment. A comparison of equivalence was then made between the sites identified from the simulation data and those from crystallographic experiments. A comprehensive list of the binding sites and their persistence across metastable states identified from SARS-CoV2, SARS-CoV and MERS-CoV M^pro^ dynamics is presented in Table 4. Sites A-L, P, R, S and V are present in all metastable states in SARS-CoV2. Based on the evolutionary conservation scores, most of the pockets (except F, J, K, N, O and U) are more conserved than the surface residues, with the strongest evolutionary signal observed for pocket B, P, R and T (Figure S10).

**Table 4:**
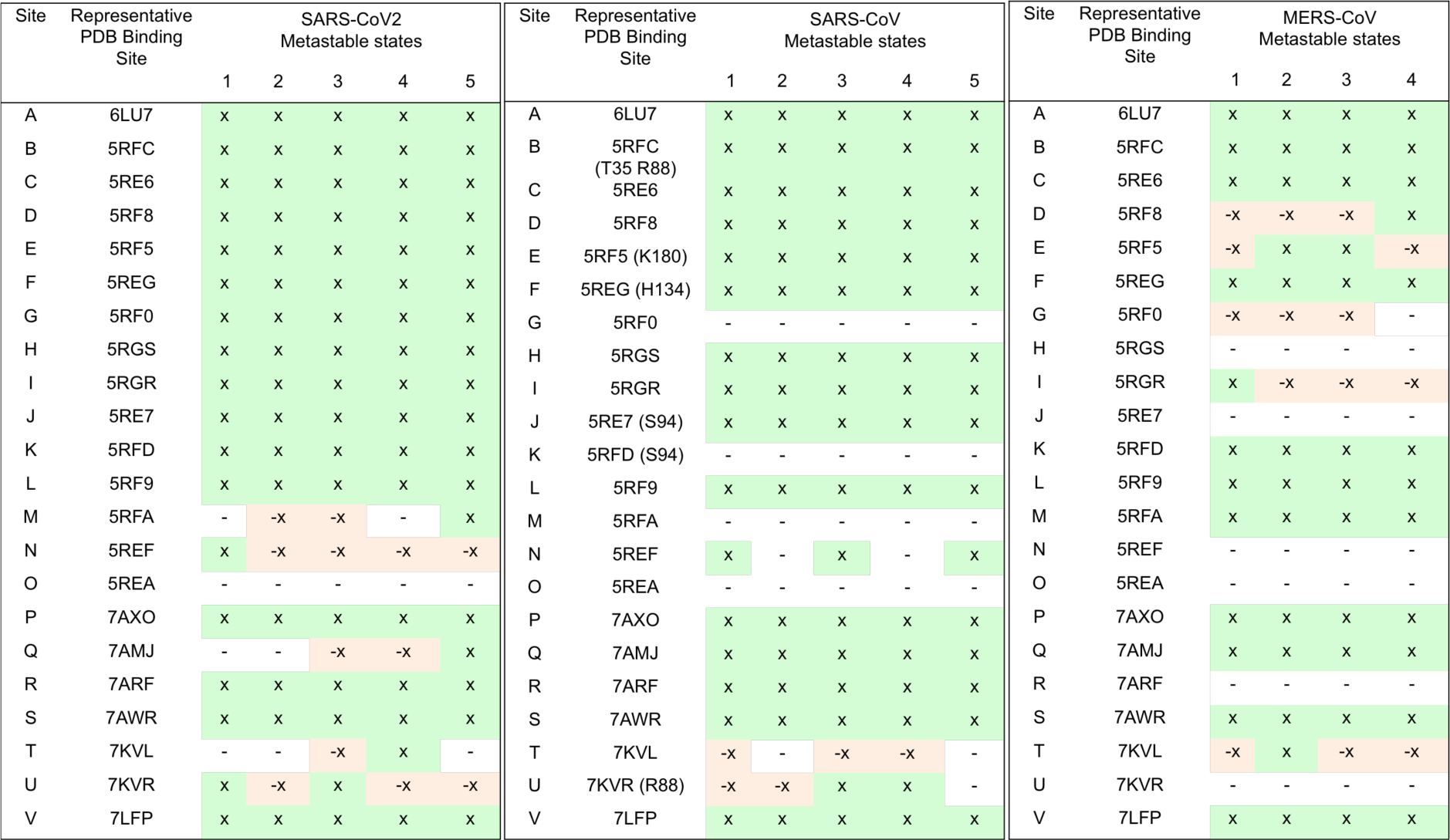
Dynamic tracking of ligand binding sites. The persistence of the ligand binding sites in (left) SARS-CoV2, (middle) SARS-CoV and (right) MERS-CoV metastable states after comparison with the representative X-ray structures. Residues in SARS-CoV that are different from SARS-CoV2 are highlighted in parenthesis. Binding sites that are present in both protomers in the metastable state and in the representative X-ray structure are indicated by a ‘x’ sign; those that are absent are noted by a ‘-’ and those that are present in at least one protomer are denoted by ‘-x’ sign.

Two copies (one in each protomer) of Site M are present in state 5, only one in states 2, 3 and none in states 1 and 4. In the crystal structure (PDB 5RFA), the carboxylic acid side chain of D295 makes interactions with the hydroxyl group side chain of T111; the side chain of Q299 makes a hydrogen bond with the backbone carbonyl oxygen atom of R4 in the N-finger; and the guanidinium side chain of R298 makes a hydrogen bond with the backbone carbonyl oxygen atom of I152. These interactions lock α-helix F in domain III to antiparallel β-barrel in domain II. The ligand occupying the large cavity at the interface further helps to stabilise the local structural elements around site M. In the absence of the ligand in the binding site and due to the dynamic fluctuations, the R298-I152 interaction is lost. The side chain of R298 is free to rotate and can adopt a conformation that can occupy the empty binding site (Figure 5A). Furthermore, the C-terminal tail also occludes one of the binding sites in state 2, and 3.

**Figure 5:**
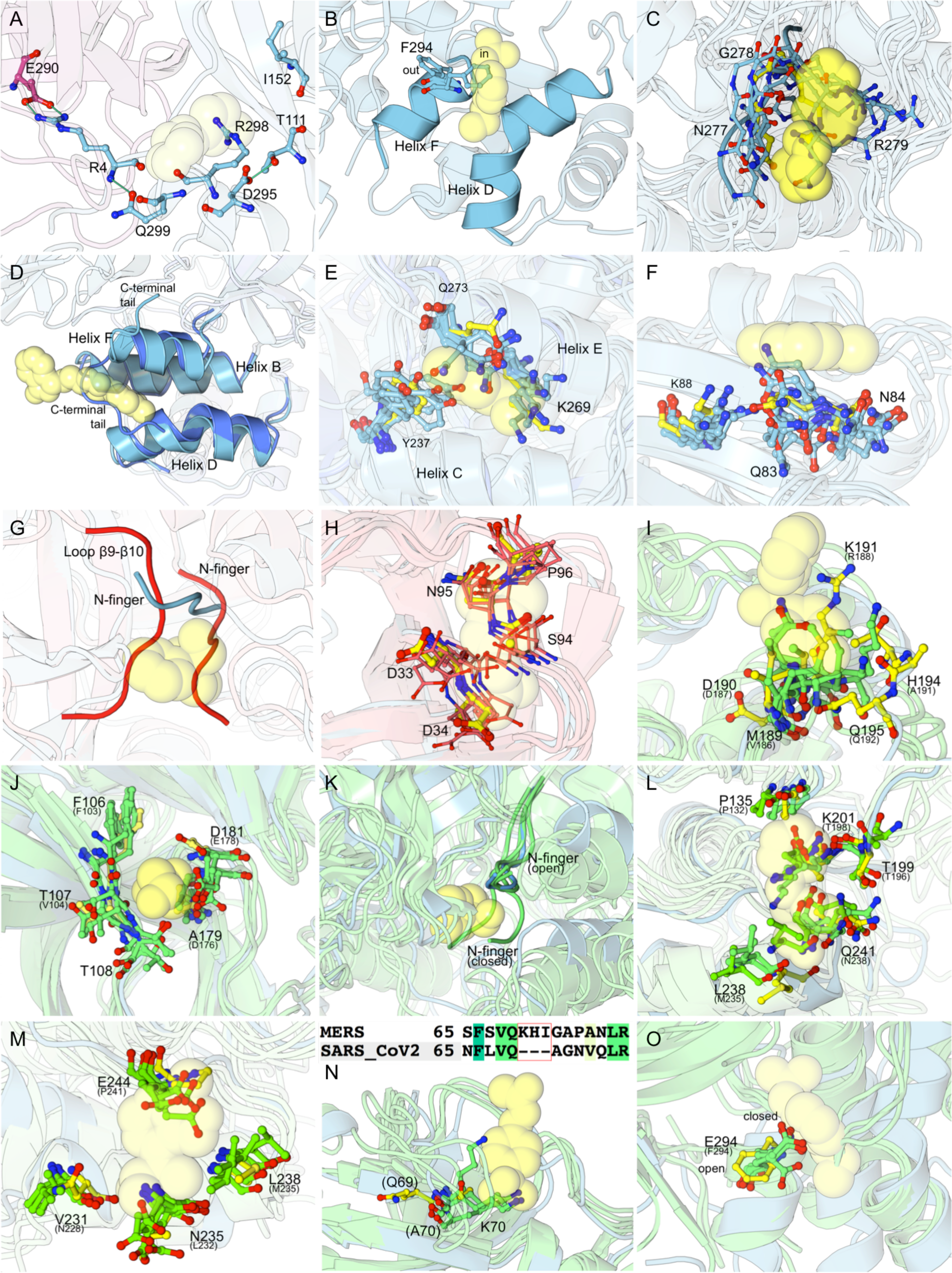
Lost ligand binding sites on (A-F) SARS-CoV2, (G-H) SARS-CoV and (J-O) MERS-CoV M^pro^. (A) Site M; when the interaction between R298-I152 is lost, the R298 side chain becomes flexible and obstructs the ligand binding site. (B) Site N; the side chain of F294 exists in two conformations. When facing inwards, it occludes the binding site. (C) Site O; is a part of a dynamic loop, which is unable to maintain the structure to which the ligand binds. Conformations of the loop from all metastable states are illustrated. (D) Site Q; in the absence of the ligand, the C-terminal tail collapses in the binding site and blocks it. Conformation of the helices from the representative PDB (id 7AMJ, cyan) and that of state 3 (dark blue) have been highlighted. (E) Site T; the side chains of Y237, K269 and Q273 (from all states) in the absence of the fragment can occupy the binding site. (F) Site U; the flexible side chains of Q83 and N84 (from all states) can disrupt the conformation on which the fragment stacks. (G) Site G; is formed at the interface when the N-finger is tucked below the substrate binding site A. Structural changes in loop β9-β10 destabilizes the N-finger, which results in its collapse on the ligand binding site. The position of the N-finger in the representative structure (PDB 5RF0) is coloured in cyan, the conformation of the state 4 is shaded in red. (H) Site K; the side chain of S94 and D33 can form a hydrogen bond, which occludes the space where the ligand binds in the representative structure (PDB 5RFD). (I) Site D; the longer side chain of M189 (in place of V186_SARS-CoV2_) obstructs the binding site. (J) Site E; a loss of steric repulsion between prevents this site to stay perpetually open; (K) Site G; the N-finger collapses on the ligand binding site as a result of the fluctuations in the β9-β10 loop. The position of the N-finger in the representative structure (PDB 5RF0) is coloured in cyan, the conformations of all metastable states are shaded in green. (L) Site H; the longer lysyl side chain of K201 (T198_SARS-CoV2_) blocks the binding site. (M) Site I; the side chain of E244 (P241_SARS-CoV2_) occludes the binding site; (N) Site J; an insertion of three residues at position 70 increases the length of the loop between β4-β5. The presence of a larger K70 side chain and the conformation of the loop restrict the dimensions of the binding site; (O) Site N; the side chain of E294 is always in the closed conformation and impedes the binding site. The representative structure is represented in cyan and yellow spheres indicate the spatial position of where the ligand binds in the corresponding representative structure.

Site N is a deep cleft between α-helix D and F and is spatially positioned adjacent to site M. The side chain of F294 (α-helix F) is shared between both the sites. The rotation of the phenyl side chain controls the opening and closure of site N. When site N is open, the phenyl side chain of F294 is positioned on α-helix F. In the closed state, the F294 side chain is positioned in the cleft. This conformation is analogous to that observed when a ligand is bound to site M. Site N is also conjoined with another larger cavity that runs orthogonal to it. When the ligand binds to this pocket (as in PDB 7AGA), the conformation of the side chain of F294 is similar to that observed in site M, which occludes site N. We observe all these conformations of F294 in our metastable states. The N site is present in both protomers in state 1; and in one of the two protomers in state 2, 3, 4 and 5. The orthogonal site is present in conjunction with site N in at least one of the protomers (Figure 5B).

Site O is a pseudo-ligand binding site on the loop between α-helix E and F. When bound, the ligand is completely solvent exposed and interacts with protein structure by making hydrogen bonds with the side chains of N277 and R279. These interactions stabilise the flexibility of this loop. In the simulated apo structure, when the ligand is absent, this loop is highly mobile and the side chains of N277 and R279 display enhanced flexibility (Figure 5C). This results in the loss of the conformation of the loop to which the ligand binds. The conformation of the loop, similar to that adopted in the representative structure is not observed in any metastable state.

Site Q at the interface between the two protomers is spatially positioned between the distal ends of α-helices B, D and F. At the end of α-helix F is a short C-terminal tail (residues 300-306). In the representative crystal structure (PDB 7AMJ), the tail orients away from the α-helical dimerization domain III and is sandwiched at the interface between domains II of both protomers, away from where the ligand binds. This provides enough space for the ligand to position in binding site Q. During the SARS-CoV2 M^pro^ apo simulations, the C-terminal tail displays dynamic flexibility and can adopt multiple conformations. Besides the conformation observed in the representative structure, one of the conformations the loop adopts occludes binding site Q and would prevent any ligand binding (Figure 5D). This conformation is similar to that observed in PDB 6LU7 structure. Site Q is present between one interface in states 3, 4; completely occluded in states 1, 2 and is present at both interfaces in state 5.

In the representative structure of site T (PDB 7KVL), the fragment makes hydrogen bonds with the hydroxyl group of Y237 (α-helix C) and the side chain of K269 (α-helix E). In the apo simulation, when the ligand is absent, the side chain of these residues can occupy the space where the fragment binds (Figure 5E). This results in the loss of this site in states 1, 2, and 5. However, the site is present in both protomers in state 4 and in only one protomer in state 3.

Site U is a solvent exposed pseudo-ligand binding site that is stabilised by the hydrogen bond interaction between the side chains of K88 and Q83. A part of this binding site is formed by N84, present in the loop between β5-β6 strands. In the absence of the fragment, the residues from this site can adopt multiple conformations, which would be unsuitable for stacking of any fragment in this site (Figure 5F). The conformation of residues comparable to the representative site is present in both protomers in states 1, 3; and in one protomer in 2, 4 and 5.

In SARS-CoV, sites equivalent to A-F, H-J, P, R, S and V are present in all metastable states. These sites are well-defined pockets and are comparable to the X-ray crystal structures of SARS-CoV2 (Table 4).

Site G, which is formed at the interface of the two protomers is lost in all metastable states of SARS-CoV. During the dynamics of the apo state, the loop between β8-β9 becomes flexible. The mobility of the loop pushes the N-finger, tucked below the substrate binding A site to collapse on site G (Figure 5G). In this conformation, no ligand would be able to bind to this site.

Binding site K is also lost in all metastable states in SARS-CoV. In this site, the hydroxyl group side chain of S94 is present (V94_SARS-CoV2_). In the absence of the ligand, the side chains of D33 and S94 orient towards each other, where they form a hydrogen bond. This stable interaction is spatially positioned on the site where the ligand binds (Figure 5H), thus completely obstructing the binding site.

Unlike in SARS-CoV2, the equivalent site on SARS-CoV, where the ligand binds in Site M is absent in all metastable states. In the representative site, the side chain of R298 forms hydrogen bond with the backbone oxygen of I152. During the simulation, this interaction is lost and the side chain of R298 in α-helix F becomes flexible and can adopt multiple conformations. One such conformation blocks the ligand-binding pocket M. The dynamics observed in this pocket are similar to that observed in SARS-CoV2 simulations (Figure 5A).

The dynamic behaviour of residues in sites N, O, T and U are also similar to that observed in SARS-CoV2 (Figure 5B/C/E/F). The formation or dissolution of site N depends upon the conformation of the phenyl side chain in F294. The site is present when the side chain orients away from the binding site and is absent when the side chain is positioned towards the binding site. Site N is observed in state 1, 3 and 5, while it is absent in state 2 and 4. Site O is a pseudo-binding site present on a highly dynamic loop. In the apo state, the N277-G278-R279 loop is highly flexible. This permits the side chains to adopt multiple conformations. However, none of the conformations are structurally similar to that which binds the ligand in the representative structure. The SARS-CoV structure lacks C-terminal tail (PDB 2C3S), hence site Q is always present in the dynamic structures. The presence of site T is depends on the conformation of Y237, Q273 and K269 side chains. In the absence of the fragment, the side chains are dynamics and can occlude the binding site. Site T is present in one protomer in states 1, 3, 4; and is absent in states 2 and 5. The dynamics of residues in site U, where R88 replaces K88, are similar to that observed in SARS-CoV2. The side chain conformation of residues on which the fragment stacks is observed in states 1, 2, 3, 4; and is absent in state 5.

From the list of 12 residues that are dissimilar between SARS-CoV2 and SARS-CoV (Figure S2) in each protomer, V35 and K88 (backbone) are present in site B. Equivalent residues in SARS-CoV are T35 and R88 respectively. These residues have similar sizes and therefore do not alter the dimensions of the binding site. However a change from V35_SARS-CoV2_ to T35_SARS-CoV_ does alter the surface charge pattern around the binding site. The side chain of K88 _SARS-CoV2_ (R88_SARS-CoV_) contributes towards stabilizing fragment binding in site U, where it makes a hydrogen bond with Q83. N180_SARS-CoV2_ is replaced with a K180_SARS-CoV_ at the entrance of binding site E. This alters the surface charge around the entrance of the binding site E towards a more positive charge. Both residues, in their respective proteins orient towards the solvent and do not interact with any other part of the protein. The backbone atoms of A94_SARS-CoV2_ (S94_SARS-CoV_) form the boundary of binding site J, while the side chain contributes to binding site K. The side chain interaction between S94 with D33 occludes the pocket and in turn has an effect on the conformation of the binding site. F134_SARS-CoV2_ is replaced with H134_SARS-CoV_ in site F. A protonated histidine side chain at the ε-nitrogen atom can form strong interactions with the ligand in SARS-CoV. V202_SARS-CoV2_, positioned at the start of helix B and is a part of the large channel-like cavity between domain II and III. Ligand AT7519 (PDB 7AGA) binds in this cavity. A deep cleft branches off this channel and forms site N. A change from V202_SARS-CoV2_ to L202_SARS-CoV_ slightly reduces the dimensions of this channel. The backbone A285_SARS-CoV2_ and the side chain of L286_SARS-CoV2_ form the boundary of the P site. A change to T285_SARS-CoV_ and I286_SARS-CoV_ does not alter the dimensions of the binding site, however these residues have been implicated in being involved in cooperative effects and enhancing dimerization in SARS-CoV ^49^. The hydroxyl side chain of S46_SARS-CoV2_ (A46_SARS-CoV_) orients near the edge of the substrate binding subsite S2. Similarly, residue 65 (N65_SARS-CoV2_ and S65_SARS-CoV_) is positioned near a cavity at the entrance of the antiparallel β-barrel in domain I, which is a potential binding site. However, we could not find any ligand that interacts with S46 or N65. Residues V_SARS-CoV2_/L_SARS-CoV_86 and S_SARS-CoV2_/A_SARS-CoV_267 are located in the core of the enzyme and do not contribute to any cavities identified on SARS-CoV2 or SARS-CoV.

In MERS-CoV, sites A-C, F, K-M, P, Q, S and V are present in all metastable states. Site D is present in both protomers in state 4 and in one protomer in state 1, 2, 3. Of particular note is the substitution of M189_MERS-CoV2_ (in place of V186_SARS-CoV2_) in this site. The longer side chain of M189 obstructs the ligand binding site in some states (Figure 5I).

Site E is present in both protomers in state 2, 3 and in one protomer in state 1 and 4. In SARS-CoV2, the side chains of D176 and E178 form the boundary of this site. In the apo state, the charge repulsion between the two negatively charges side chain prevents the closure of this site in SARS-CoV2. However, D176_SARS-CoV2_ is replaced with A179_MERS-CoV_ and E178_SARS-CoV2_ with D181_MERS-CoV_. In the absence of the ligand, and with no charge repulsion between the negatively charged side chains, the side chain of D181 obstructs the binding site in some metastable states (Figure 5J).

Site G is present in one protomer in states 1, 2, 3 and is absent in state 4. In SARS-CoV2, the N-finger is tucked below site A, which provides enough space at the interface for the ligand to bind in site G. In the simulated apo state of MERS-CoV and similar to that observed in SARS-CoV2, the N-finger can also collapse and occupy the binding site resulting in its closure (Figure 5K).

In Site I, N228_SARS-CoV2_, L232_SARS-CoV2_, M235_SARS-CoV2_ and P241_SARS-CoV2_ are replaced with V231_MERS-CoV2_, N235_MERS-CoV2_, L238_MERS-CoV2_ and E244_MERS-CoV2_ respectively. The longer carboxylic side chain in E244_MERS-CoV_ can adopt a conformation that obstructs the binding site (Figure 5L). This is observed in at least one protomer in state 2, 3 and 4; while the binding site is clear in state 1.

Site T is present in both protomers in state 2; and in one protomer in states 1, 3, 4. Here, Y273_MERS-CoV_ in substituted in place of L275_SARS-CoV2_. Furthermore, a large indole ring in W236_MERS-CoV_ replaces the smaller side chain of V233_SARS-CoV2_, making the binding site shallow than its representative structure. Taken together, the side chains of W236_MERS-CoV_ and Y273_MERS-CoV_ act like a wedge to split and widen α-helices C and E. Therefore, site T is persistently more open when compared with the dynamics of SARS-CoV2 or SARS-CoV.

Sites H, J, N, O, R and U are absent in all metastable states in MERS-CoV. In site H, the substitution of a shorter hydroxyl group in T198_SARS-CoV2_ to a longer lysyl side chain in K201_MERS-CoV_ completely obstructs the binding site in all metastable states (Figure 5M). In MERS-CoV, loop β4-β5 is extended by insertion of three residues between positions 69-70. As a result there is a change from A70_SARS-CoV2_ to a lysine at this position. The longer lysyl side chain obstructs site J where the ligand binds (Figure 5N). F294_SARS-CoV2_ is substituted with E294_MERS-CoV_ in site N. Unlike in SARS-CoV2 and SARS-CoV, the side chain of E294_MERS-CoV_ point towards the N site cleft, which blocks site N (Figure 5O). Ligands interact with site S by forming a disulphide bond with C156_SARS-CoV2_. However, in MERS-CoV, the cysteine residue is replaced with V159_MERS-CoV_, which would prevent any disulphide bond formation. In site U, the side chain on which the ligand stacks is absent due to the substitution of N84_SARS-CoV2_ by G87_MERS-CoV2_.

## Discussion

Despite tremendous advances in the inhibitor design for SARS-CoV2 M^pro^ enzymes, our understanding of the role of structural dynamics of the experimentally identified ligand binding sites remain largely uncharacterized. Most molecular dynamics studies have focused only on the substrate binding site of the M^pro^ enzyme.^50–52^ Other computational studies have looked into identifying novel pockets and investigating allostery.^53, 54^ However, these studies are limited in comparing dynamics with the vast crystallographic data available on ortho- and allosteric ligand binding sites across β-coronavirus homologs.

In this study, we map all non-redundant ligand-binding sites reported in the PDB for β-coronavirus M^pro^ enzyme homologs including SARS-CoV2, SARS-CoV and MERS-CoV. We perform 25 µs MSM-based adaptive sampling MD simulations to study the dynamics of the binding sites. It is worth noting that we simulated the apo form of the SARS-CoV2, which was generated by the removal of the ligand from the substrate-binding site in PDB 6LU7. However, this does not have any impact on our analysis as we sample all crystallographic conformations. The analysis emphasizes that even though the β-coronavirus M^pro^ structures are very similar, they display remarkable structural dynamics. The differences in dynamics are subtle and indistinguishable using conventional methods. We therefore employed dynamically sensitive CVAE-based machine learning approaches to resolve the differences between each system. MSMs were built to identify kinetically relevant metastable states, which were then used to study the spatiotemporal evolution of the ligand binding sites. The metastable states generated from the simulations were searched for the presence of pockets and compared individually with all other experimentally derived crystal structures representing non-redundant ligand binding sites.

The M^pro^ enzymes are homodimers and each binding site is present as two copies, one on each protomer except for site V. The dynamical behavior of each protomer is stochastic and independent of the other. This is evident from the structural dynamics of the binding sites, which in some metastable states appear only in one protomer and absent in the other. Our finding is supported by previous work on M^pro^ enzymes where the dynamics of different protomers map on the different regions of conformational space.^50^ We also identify that loops connecting different structural features are the most flexible regions of the enzyme and contribute towards the local motions, while movement between the two coaxially stacked protomers contribute to the global dynamics. The presence or absence of binding sites in each protomer is independent of the influence of the adjacent protomer except for the sites at the interface. The ligands that bind at the interface work by stabilizing the global motions that contributes towards inhibiting mechanistic function.

To assess the possibility of a broad-spectrum inhibition of M^pro^ enzymes, we analyzed the structural and dynamic conservation of the binding sites across the three β-coronavirus homologs. We rationalized that an inhibitor designed to target a conserved binding site would have relatable effects across homologs. This would be advantageous for the design of therapeutics in dealing with any future viral outbreaks. We analyzed the dynamics of the ligand binding sites by comparing the sequence and structural features between relative homologs.

SARS-CoV2 and SARS-CoV have 96% similar sequence identity. We identify that of the 12 residues (out of 306) that are different between SARS-CoV2 and SARS-CoV (Figure S2) in each protomer, 8 are associated with an experimentally identified ligand-binding site. The substitution of some of these residues have an effect on the surface charge pattern (N180_SARS-CoV2_/K180_SARS-CoV_ and T35_SARS-CoV2_/V35_SARS-CoV_), interactions (F134_SARS-CoV2_/H134_SARS-CoV_) dimensions (V202_SARS-CoV2_/L202_SARS-CoV_), enhancing enzymatic activity via dimerization (A285_SARS-CoV2_/T285_SARS-CoV_ and L286_SARS-CoV2_/I286_SARS-CoV_) or completely block the space where the ligand binds (A94_SARS-CoV2_/S94_SARS-CoV_). One substitution (K88_SARS-CoV2_/R88_SARS-CoV_), has no notable effect on the binding site. 2 residues (S46_SARS-CoV2_/A46_SARS-CoV_ and N65_SARS-CoV2_/S65_SARS-CoV_) are a part of potential cavities but no ligand has been identified to bind to them yet. The remaining 2 residues (V86_SARS-CoV2_/L86_SARS-CoV_ and S267_SARS-CoV2_/A267_SARS-CoV_) are located in the core of the enzyme and are not solvent accessible.

We then tracked the dynamic persistence of the ligand binding sites in the MSM-derived metastable states in the three homologs and made comparisons with the representative binding sites from the crystal structures. All of the identified binding sites are located on the surface of the M^pro^. Ligand binding sites A-L, P, R, S, V (SARS-CoV2); A-F, H-J, L, P-S, V (SARS-CoV); and A-C, F, K-M, P, Q, S and V (MERS-CoV) are present in all metastable states. Site O is the only ligand binding site that is absent in all homologs. Site O is a pseudo-binding site on a solvent exposed loop whose conformation once lost is never observed in the dynamics of apo M^pro^. Sites M, N, Q, T, U in SARS-CoV2; N, T, U in SARS-CoV; and D, E, G, I, T in MERS-CoV are present in some states and absent in others. Sites G, K, M, O (SARS-CoV) and H, J, N, O, R and U (MERS-CoV) are completely absent in their respective homologs. It is worth noting that there are multiple binding sites that lie adjacent to one another e.g. sites B and C; P and T; R and S. Fragments occupying these sites can be chemically linked to enhance effective binding (Figure S11). Furthermore, there are several other structural features present around the experimentally identified binding sites, which can be exploited to improve the design of inhibitors. For example, empty cavities are present adjacent to sites H, K, L, N, Q and S (Figure S12). These cavities can be used as extensions of existing binding sites to improve ligand design.

Our detailed structural dynamics analysis highlights the importance of the dynamic conservation of ligand binding sites across β-coronavirus homologs. Based on these observations we emphasize that ligand design should be preferred on target binding sites that are not only structural but also dynamically conserved across all β-coronavirus homologs.

## Conclusions

The past 20 years has seen outbreaks caused by three highly pathogenic β-coronavirus namely SARS-CoV in 2002, MERS-CoV in 2013 and SARS-CoV2 in 2019.^55^ The social and economic impact of the current pandemic has been exceptional. This crisis has led to an urgent requirement to develop therapeutics. Even though a number of vaccines have been approved by the Food and Drug Administration, alternative strategies targeting essential viral components are required as a back-up against the emergence of lethal viral variants. One such target is the main protease that plays an indispensible role in viral replication.^18, 19^ Multi-nodal, large interdisciplinary consortiums have reported potential drug candidates.^37, 39, 56^ The availability of M^pro^ X-ray structures in complex with inhibitors provides unique insights into ligand interactions. This data in conjunction with molecular simulations can aid to further improve design of inhibitors including exploring the dynamic conservation of ligand binding sites across β-coronavirus homologs that are highly relevant to human disease. Employing such a strategy is essential in preparing towards any future viral outbreaks.

## Experimental Methods

### Ligand binding site identification

The protein data bank in Europe knowledge base (PDBe-KB) was searched with the key word “3C-like proteinase” and selecting “Severe acute respiratory syndrome coronavirus 2 (2019-nCoV)” as the organism. The PDB codes were noted and the structural coordinates downloaded. Thorough analysis was done by superimposition of the structures. The key interacting residues were identified within a 4.0 Å cut-off distance around the ligand. This was repeated until all entries were evaluated. From this list, a non-redundant representative structure for each binding site was identified. For example in the PDB 6LU7,^22^ the ligand N3 interacts with residues C145, H41, G189, P168, E166, H163 and H164 in the substrate binding site. Thus, 6LU7 was selected as the representative structure for all ligands that interacted with these residues and labelled ‘site A’. Figures for representative structure and ligands were generated using Protein Imager.^57^

A similar protocol was applied for SARS-CoV and MERS-CoV M^pro^ structures and non-redundant representative structures were identified. PDB identifiers, structural analysis and ligand interaction data are listed in the supplementary section. The non-redundant representative ligand binding site data has been tabulated in Table 1-3.

### Adaptive Sampling molecular dynamics simulations

The coordinates of the apo structure of the SARS-CoV2 (PDB 6LU7),^22^ SARS-CoV (PDB 2C3S),^58^ and MERS-CoV (PDB 4YLU)^59^ protease in their dimeric form were downloaded to run molecular dynamics (MD) simulations. Ligands and all crystallisation agents/additives were removed from their respective binding sites. The protonation state of all titratable side chains were determined using *ProteinPrepare* functionality as implemented in HTMD framework.^60, 61^ The charges were assigned after optimisation of the hydrogen-bonding network in the protonated structure.^61^ The catalytic cysteine residue was set to a reduced state. The Amber ff14SB force field was used to describe the protein.^62^ Each system was solvated using TIP3P water in a cubic box, the edge of which was set to at least 10 Å from the closest solute atom.^63^ Counter ions were added to neutralise the system. The simulation protocol was identical for each system. The systems were minimized and relaxed under NPT conditions for 50 ns at 1 atm. The temperature was increased to 300 K using a time step of 4 fs, rigid bonds, cut off of 9.0 Å and particle mesh Ewald summations switched on for long-range electrostatics.^64^ During the equilibration step, the protein’s backbone were restrained by a spring constant set at 1 kcal mol^-1^ Å^-2^, while the ions and solvent were free to move. The production simulations were run in the NVT ensemble using a Langevin thermostat with a damping constant of 0.1 ps and hydrogen mass repartitioning scheme to achieve a time step of 4 fs.^65^ The final production step was run as Adaptive Sampling, without any restraints, as multiple iterations of short parallel simulations as implemented in HTMD framework.^60^ Each system was run for 125 epochs (iterations) and each epoch consists of four parallel simulations of 50 ns each, equalling 25 µs of simulated time. The short simulations after each epoch are postprocessed based on the backbone dihedral angle metric. A rough Markov model is then used to decide from which part of the configuration space to respawn the following simulations in the next epoch. Visualization of the simulations was done using the VMD package.^66^

### Markov state models

Markov state models (MSMs) were constructed to provide kinetics and free energy estimates. The MSM was built using the PyEMMA v2.5.7 program.^67^ It was not possible to build an MSM using just the features of the 24 dissimilar residues (12 in each protomer) between SARS-CoV2 and SARS-CoV. Therefore, all backbone dihedral angles were selected. In addition, the first χ angle (χ1) from 24 dissimilar residues were also included in MSM building. For MERS-CoV, χ1 angles from residues at equivalent position were also selected. Time-lagged independent component analysis (tICA) was used to reduce the dimensionality of the data.^68, 69^ It was possible to build models that were Markovian with a lag time of ≥10, with the lag time being selected according to the convergence of the implied timescales. The dimension reduction was achieved by projecting on the three slowest tICA components. The *K-means* clustering algorithm was used to obtain 100 microstates. The conformational clusters were grouped together based on kinetic similarity using the PCCA+ algorithm.^70^ The PCCA+ algorithm uses the eigenvectors of the MSMs to group together clusters, which are kinetically close, resulting in a set of macrostates. The final number of metastable macrostates was selected based on the implied timescale plot. The MSM were validated using Chapman-Kolmogorov test implemented in PyEMMA.^67^

### CVAE-based Deep learning implementation

The Convolutional Variational Autoencoder or CVAE was used for analysis,^40^ which has been optimized for large scale systems on HPC platform.^71^ The implementation of CVAE has been previously shown to provide meaningful insights to diverse systems such as protein folding,^72^ enzyme dynamics,^41, 73^ Coronavirus spike protein ^74^ and Coronavirus non-structured proteins.^75^

A CVAE consists of a variational autoencoder along with multiple convolutional layers. Generally, the autoencoder (AE) has an hourglass type of shape where high dimensional data goes into as input and the AE captures only the essential information required to represent the original input data. This compressed latent representation is then used to reconstruct the data back to the original format ensuring no loss of information during the compression phase. The variational approach at the latent space is included as an additional optimization requirement. The introduction of variational technique forces the compressed key information to normally distribute over the latent space. Convolutional layers are used instead of feed forward layers because the convolutional layers are more effective at detecting and captureing both the local and global patterns in the input data especially where the data has multi-layered structures like complex proteins as presented here. The complete CVAE structure is shown in Figure 6A with different steps that are performed from raw simulation data to resolution of β-coronavirus M^pro^ solely based on their local and global conformational dynamics.

**Figure 6:**
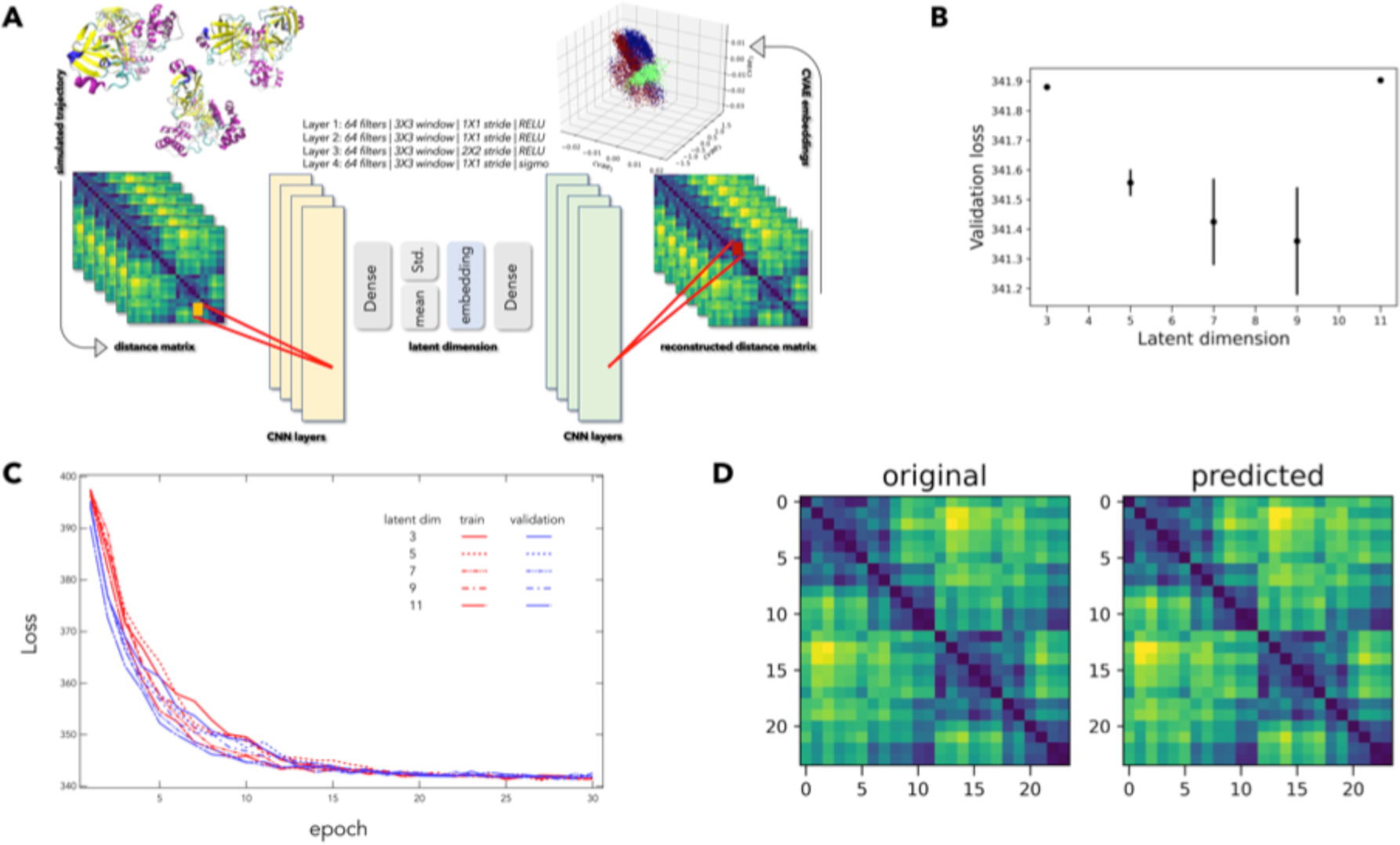
CVAE based Deep Learning Implementation. (A) Complete CVAE architecture where distance matrix is used as input data to feed into the CVAE for generating low-dimensional representation. The input data is then trained where the training quality can be followed by training and validation loss at different dimension over the epochs. (B) The validation loss is plotted at different latent dimension to determine optimum values of the low dimension. (C) Simultaneously the training and validation loss is assessed over consecutive epochs at various dimensions. (D) A comparison between original input and predicted data to ensure no loss of information during compression process.

The distance matrix of the 24 x 24 dissimilar Cα atoms was used as input for the CVAE architecture. Using the Horovod library, the data parallel model was trained on the Summit supercomputer. Each CVAE was trained for a fixed number of epochs based on the convergence of loss and variance-bias trade-off. Each training utilized up to 16 Summit nodes (96 V100 GPUs), and the effective batch size being the sum of every individual training instance. Therefore, the individual batch size was selected to be relatively small to avoid the generalization gap for large-batch training. The dataset was divided into training/validation (80:20 % of the simulation trajectories) and randomly shuffled. To search for the optimal clustering and reconstruction quality of the CVAE, the training procedure was repeated for various latent dimension sizes and to identify the best model for the dataset (Figure 6B). The loss over the epochs is as expected (i.e., without over fitting or any other unusual behavior) and shown in Figure 6C. Finally, the original input data was compared with the predicted (i.e., decompressed) data to ensure no loss of information during the compression process through the latent space (Figure 6D).

### Dynamic pocket tracking

Pocketron was used to detect small molecule binding sites using default values.^76^ The metastable states were screened for pockets, which were classified as open if they could accommodate at least 5 water molecules (coarse equivalent of a small fragment). Each representative binding pocket, identified from the crystal structures, was compared by superimposition with the metastable state from each system.

### Analysis of pairwise correlated positions in evolution

Pairwise evolutionary constraints were estimated from a multiple sequence alignment (MSA). The FASTA sequence from the SARS-CoV2 M^pro^ (PDB 6LU7) was selected as reference and the MSA was built using hhsuite3.^77^ Pairwise correlations were calculated using ccmpred package ^78^ as per the parameters described in Akere et al.^73^ Raw correlation scores (C_i_) were then scaled as per Kamisetty et al.^79^ For all 22 pockets (see Table 1), the scaled pairwise correlation matrix was used to estimate the evolutionary conservation score (E_a_) of each pocket (Eq 1), where N is the number of residues in the pocket.

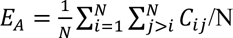

The score estimates the evolutionary constraints on the pocket as an average of the pairwise correlation in the pocket. For reference, scores were compared with the median and standard deviation of C_i_ for all surface residue pairs (Figure S10). Surface residues were defined as having > 50% relative accessible surface area.^80, 81^

## Acknowledgements

The authors would like to thank the UK High-End Computing Consortium for Biomolecular Simulation, HECBioSim (http://hecbiosim.ac.uk) for time to run simulations on ARCHER. BI would like to acknowledge the COVID-19 pump-priming grant from the University of Huddersfield for funding computing resources for analysis. SH would like to thank Prof Frank Kozielski for insightful discussions on the manuscript.

This material is based upon work supported by the U.S. Department of Energy, Office of Science, through the Advanced Scientific Computing Research (ASCR), under contract number DEAC05-00OR22725 and the Exascale Computing Project (ECP) (17-SC-20-SC). This work was performed at the Oak Ridge Leadership Computing Facility (OLCF) of the Oak Ridge National Laboratory (ORNL) and used the Extreme Science and Engineering Discovery Environment (XSEDE) ^82^ COVID-19HPC Consortium at the IBM AC922 Summit supercomputer of the OLCF at ORNL through allocation TG-ASC200020. The ASCR, the ECP and the National Virtual Biotechnology Laboratory (NVBL) help in implementation of deep learning algorithms, data processing and analysis respectively.

## Data Availability Statement

The trajectories of M^pro^ simulations and models of the metastable states can be obtained from the corresponding author. Jupyter-notebooks to generate Markov State Models can be downloaded from 10.6084/m9.figshare.14343725

## Author Contributions

Data mining and collation: EC, MR, RA, SM, MS, MM, KB, BI; Binding site analysis: EC, KB, BI, SH; Co-evolution analysis: SD, AP; DL: HM, AR, DB, JY, SH; Simulations: SH, HM, AR, BI; MSM: SH, AM; Manuscript writing: EC, AR, DB, SH; other inputs: all co-authors

## Funding

The authors declare no potential conflict of interest

## Conflict of Interest

The authors declare no potential conflict of interest

## Supplementary Material

The supporting information is available free of charge at..

Mapping the binding sites; Sequence alignment between SARS-CoV2, SARS-CoV and MERS-CoV M^pro^; Superimposition of SARS-CoV2 and SARS-CoV M^pro^ structures; Interactions between SARS-CoV2 and ligands in their representative ligand binding sites; Interactions between the SARS-CoV2 M^pro^ in the pseudo-ligand binding sites; Conformational drift in M^pro^ enzymes; Root mean squared fluctuation plots of M^pro^ enzymes; Markov State Model of SARS-CoV2, SARS-CoV and MERS-CoV M^pro^ enzymes; Evolution conservation score for each pocket; Adjacent binding sites in SARS-CoV2; Multiple ligands binding in sites. (PDF)

Appendix Tables S1-S6: Details of the ligand binding sites (PDF)

## Notes

### Competing Interest Statement

The authors have declared no competing interest.

### Summary of Updates

Supplemental files updated

