## Supplementary information for "Structural dynamics of the β-coronavirus M^pro^ protease ligand binding sites"

### Table of Contents


Mapping the binding sites 2

**Figure S1.** Sequence alignment between SARS-CoV2, SARS-CoV and MERS-CoV $M^{pro}$. 8

**Figure S2**. Superimposition of SARS-CoV2 and SARS-CoV $M^{pro}$ structures. 9

**Figure S3**. Interactions between SARS-CoV2 and ligands in their representative ligand binding sites. 10

**Figure S4.** Interactions between the SARS-CoV2 $M^{pro}$ in the pseudo-ligand binding sites. 11

**Figure S5.** Conformational drift in $M^{pro}$ enzymes. 12

**Figure S6.** Root mean squared fluctuation plots of $M^{pro}$ enzymes. 13

**Figure S7.** Markov State Model of SARS-CoV2 $M^{pro}$ enzyme. 14

**Figure S8**. Markov State Model of SARS-CoV $M^{pro}$ enzyme. 15

**Figure S9**. Markov State Model of MERS-CoV $M^{pro}$ enzyme. 16

**Figure S10.** Evolution conservation score for each pocket. 17

**Figure S11.** Adjacent binding sites in SARS-CoV2. 18

**Figure S12**. Multiple ligands binding in sites. 19

#### Mapping the binding sites

Most of the ligands that have been crystallised bind to the substrate binding site (site A in Figure 2A). We identified 185 ligands that bind either covalently or non-covalently, exploiting various subsites of this pocket. The substrate binding site is a deep cleft between domains I and II. C145 is spatially positioned in the deep distal end of the pocket. For inhibitors to covalently attach to C145, they would have to penetrate deep inside the binding site. The non-covalent inhibitors occupy one of the subsites within the substrate binding site. PDB 6LU7 was chosen as the representative structure to highlight this binding pocket (Jin et al., 2020a).

Sites B (PDB 5RFC; (Douangamath et al., 2020)) and C (PDB 5RE6; (Douangamath et al., 2020)) are present in domain I and form part of the antiparallel β-barrel structure (Figure 2B/C). The two sites lie adjacent to one another, separated by a part of the β4 strand (I78-M82). The N- group in methyl (2-methyl-4-phenyl-1,3-thiazol-5-yl)carbamate interacts with the backbone carbonyl oxygen of H80 and the pyrimidine group in N-{4-[(pyrimidin-2-yl)oxy]phenyl}acetamide in site C is positioned above H80. Five ligands have been co-crystallised in site B, while two have been identified to bind in site C.

Site D (PDB 5RF8; (Douangamath et al., 2020)) is a shallow, surface exposed binding site formed as a part of a large hairpin loop that connects domain II and III (residues P184-Q192) (Figure 2D). 4-amino-N-(pyridin-2-yl)benzenesulfonamide (Z271004858) is anchored in this binding site via hydrogen bonds with R188 on one end and backbone carbonyl oxygen of A191 on the other (Figure S2D). It is noteworthy that site D shares residues R188-Q192 with the substrate binding site (site A). Comparing the conformation of the linker loop that connects domain II with III in site D, with the structure of the loop when the ligands are present in their respective binding sites in site A, suggests that only one binding site can be occupied at a time. Ligand binding in site A shifts the conformation of the guanidinium side chain of R188 such that binding site D is lost. However, ligand binding to site D has no effect on subsite S4 of the substrate binding pocket. Only one ligand has been reported to bind in site D.

Site E (PDB 5RF5; (Douangamath et al., 2020)) is a pocket formed between the β7-β8 loop and the proximal end of the linker loop in domain II (Figure 2E). The pocket is so small that it can accommodate only a small fragment. Larger fragments also bind to this site by anchoring inside this pocket. Four fragments have been identified that bind to this pocket including 1,1 bis(oxidanylidene)thietan-3-ol (Z3241250482) that has been used as its representative structure.

Site F (PDB 5REG; (Douangamath et al., 2020)) is formed from residues from three different loops including β1-β2 (R105, Q107), β3-β4 (F134) in domain II and the linker loop (G183). The binding site is small and can only accommodate half of the (2~{S})-~{N}-(4-aminocarbonylphenyl)oxolane-2-carboxamide (Z1545313172), with the oxolane-carboxamide group inside the binding pocket and the aminocarbonylphenyl group on the outside. The backbone of G183 makes hydrogen bond and F134 interacts with the ligand by forming π-stacking interactions. The binding pocket is also connected to another, larger pocket via a narrow passage. However, the passage is not wide enough for a phenyl group to pass through it (Figure 2F).

Site G (PDB 5RF0; (Douangamath et al., 2020)) is a pocket, present just below the substrate-binding site (site A), at the interface between the two protomers. The site is sandwiched between K137 and G138 (protomer A) and the N-finger residues (S1-A7) of protomer B. A part of this binding site is also formed by a loop between β11-β12, which makes the lower lip of binding site A (L166-V171). L282, G283 and N214 from the α-helix B in domain III also contribute to this pocket (Figure 2G). Binding of a ligand in the substrate-binding pocket (site A) alters the conformation of the β11-β12 loop at the proximal end of the binding site near S4 subsite. This change in conformation of the loop has an effect on the dimensions of site G. Only one ligand POB0073 ([1-(pyridin-2-yl)cyclopentyl]methanol) has been identified that binds to this pocket.

Site H (PDB 5RGS; (Douangamath et al., 2020)) is present in the α-helical dimerization domain III (Figure 2H). The site is formed from two subpockets that are separated by the side chains of T198 and E240. Residues form β7-β8 loop (P132, N133) from domain II, the linker loop (G195, T196, T198), the loop between α-helix C and D in domain III (N238, Y239, E240) and A234, M235 from the terminal end of α-helix C form this binding site. The methanol group makes hydrogen bonds with the carbonyl oxygen atom from the backbone of M235 and the morpholino oxygen makes interactions with the side chain of N238. Four ligands have been identified that can bind to this pocket, while Z1259086950 has been used as a representative ligand to describe this site.

In the representative structure for Site I (PDB 5RGR; (Douangamath et al., 2020)), the fragment N,1-dimethyl-N-(propan-2-yl)-1H-pyrazolo[3,4-d]pyrimidin-4-amine (Z328695024) binds at two sites on $M^{pro}$. The first site is the same site as site E, where the N-propanyl group is tucked inside the pocket and the pyrazolo-pyrimidine stacks on the β7 strand of the 2nd antiparallel β-barrel in domain II. The second site (I2) is formed by residues from α-helix C (N228, L232, M235) and the loop between α-helix C and D (P241) in the dimerization domain III (Figure 2I). The binding site is an exposed, shallow pocket. Only one ligand has been identified to bind to this site.

Site J (PDB 5RE7; (Douangamath et al., 2020)) is formed on the distal end, at the entrance of the antiparallel β-barrel in domain I. This site is a part of a long channel that runs between domains I and II (Figure 2J). Residues from β1 (W31), loop β4-β5 (A70, G71), β5 (V73, L75), and the loop between β6 from domain I and β7 from domain II (T93, A94, P96, K97) lie within 4 Å from the ligand. The N-sulfamoyl group and the phenyl ring in N-[(4-sulfamoylphenyl)methyl]acetamide (Z30932204) are positioned in the site while the acetamide group protrudes into the solvent.

Site K (PDB 5RFD; (Douangamath et al., 2020)) is a shallow pocket formed between the loops that connect β2 and β3 in antiparallel β-barrel in domain I and the loop that connects domains I and II (Figure 2K). It lies is adjacent to site J and residues A94, N95, P96 (β6-β7 loop) are shared between the two sites. The ligand 2-[(methylsulfonyl)methyl]-1H-benzimidazole (Z126932614) in site K is solvent exposed. Ligand binding to site K has no effect on the dimensions of site J. Three ligands have been identified that are bound to site K.

Site L (PDB 5RF9; (Douangamath et al., 2020)) is a long, elongated channel like site located immediately downstream from site K, along the loop that connects domain I and II (residues P96-K100). One end of the site is a deep cavity formed from K12, K100, D155, G156, V157 and binds the pyrazol ring in 1-[(2~{S})-2-methylmorpholin-4-yl]-2-pyrazol-1-yl-ethanone (Z217038356). The ketone oxygen anchors the inhibitor in the binding site by making interactions with K12. The methyl group substituted of the morpholino ring projects away from the binding site towards site K (Figure 2L). Three ligands have been identified to bind in this site.

Site M (PDB 5RFA; (Douangamath et al., 2020)) is a deep cavity formed at the interface between the two protomers. The site is formed between domain II and III on protomer A, including residues from the N-terminal (M6, A7, F8, P9), α-helix F (D295, R298, Q299), β8 (T111, S113) and β9 (Q127). Residues Y118 (loop β8-β9), S123, G124 (β9) and L141 (loop β9-β10) from protomer B contribute to this site. When 1-methyl-N-{[(2S)-oxolan-2-yl]methyl}-1H-pyrazole-3-carboxamide (Z2643472210) binds, the pyrazole ring penetrates the binding site between the N- and C-terminal, while the amide group and the methyl oxolane ring interacts with residues from protomer B. The amide group makes hydrogen bond with the backbone carbonyl oxygen of S123 from protomer B. Two ligands have been identified that bind to this site.

Site N (PDB 5REF; (Douangamath et al., 2020)) is sandwiched between α-helix D and F. The site forms a deep cleft that branch off an extended channel of cavities formed between the antiparallel β-barrel in domain II and the α-helices in dimerization domain III (Figure 2N). The methylsulfonylamino group in methyl 3-(methylsulfonylamino)benzoate (Z24758179) faces the channel, while the benzoate group is sandwiched between F294 and I249. Residues from α-helix F are shared between Site M and Site N. Ligand binding to site M alters the conformation of the side chain of F294, which completely occludes site N. When the ligand is bound in site N, the phenyl side chain rotates away from the binding site and is positioned above α-helix F, providing enough space for the ligand to bind. Another ligand, AT7519 (PDB 7AGA; (Günther et al., 2020)) also binds to site N, not in the cleft but in an orientation that is along the channel. When this happens, the side chain of F294 adopts a similar conformation when a ligand is bound to site M, which obstructs the deep cleft.

Site O (PDB 5REA; (Douangamath et al., 2020)) is a solvent exposed, pseudo-ligand binding site where (azepan-1-yl)(2H-1,3-benzodioxol-5-yl)methanone (Z31432226) forms interactions with N277, G278 and R279 (Figure 2O). These residues are part of the loop between α-helices E and F. There is no cavity in which the ligand binds but is anchored to the exterior of $M^{pro}$ via hydrogen bonds with the side chain of N277 and R279. The ligand in site O is positioned in site J, in its symmetry-related protomer (Figure S4).

Site P (PDB 7AXO; (Günther et al., 2020)) is a pocket formed by residues at distal ends of α-helices C and E and the loop that leads to α-helix F, in the dimerization domain (Figure 2P). The phenyl ring in N-hydroxy-4-{[(2S)-3-methyl-2-phenylbutanoyl]amino}benzamide (AR-42) is tucked in a hydrophobic subpocket formed by residues M276, A285, L276 and L287. The central amide anchors the ligand to the backbone carbonyl oxygen of L272, while the benzamide group is exposed to the solvent (Günther et al., 2020). Two ligands have been identified that bind to this pocket.

Site Q (PDB 7AMJ; (Günther et al., 2020)) is formed at the interface of the dimerization domains, with contribution from the residues at the distal ends of α-helices B, D and F in domain III in one protomer and the loop between β9 and β10 (residues L141, N142) in the other protomer (Figure 2Q). It is worth noting that N142 also forms a part of the substrate binding site (site A). The piperazine ring and its dimethylphenyl ring substituent in PD168568 occupy a hydrophobic pocket formed from the three-helix bundle present in the dimerization domain. The isoindolone group interacts with N142 at the entrance of the substrate binding site in the adjacent protomer. Five ligands have been identified that interact with this binding site.

Site R (PDB 7ARF; (Günther et al., 2020)) is present in domain II. Residues from β7 (K100, Y101, K102) and β11 (D155, C156) contribute to this shallow, solvent exposed binding site (Figure 2R). Small fragments bind to this site by covalently attaching to the surface exposed C156 via a disulphide bond. Fragments like thioglucose, can make additional interactions with the backbone carbonyl oxygen of Y101 and K102. Six fragments have been identified that covalently bind in this binding site.

Site S (PDB 7AWR; (Günther et al., 2020)) is a shallow, surface binding pocket between domain I and II, formed by residues D33, T98-F103 (Figure 2S). Tegafur binds to the site via π-stacking interactions between Y101 and the pyrimidine ring of the ligand. Site R and S are separated by common residues from the β7 strand. A glutathione isopropyl ester (PDB 7AX6) straddles β7 strand and interacts in both site R and S simultaneously.

Site T (PDB 7KVL; (Noske et al., to be published)) is sandwiched between α-helices C (V233, Y237) and E (K269, Q273) in the dimerization domain (Figure 2T). The nitrogen in the pyridine ring makes interactions with the hydroxyl group in the Y237 and the K269 side chain forms a hydrogen bond with oxygen in the carboxamide group of the fragment. This site lies adjacent to site P and is separated by the side chain of Y237.

Site U (PDB 7KVR; (Noske et al., to be published)) is a pseudo-ligand binding site formed at the interface of two symmetry-related protomers (Figure 2U). The dimethylpyridine-2,4-diamine fragment stacks on side chains from residues in the loop between β5-β6 strands (Q83, N84). A hydrogen bond between the side chains of Q83 and K88 (β6 strand) further stabilizes this stacking platform. A water molecule bridges interactions between the fragment and the backbone oxygen atom of M82. The fragment also makes hydrogen bond with the side chain of D48 of the symmetry-related protomer (Figure S4).

Site V (PDB 7LFP; (Noske et al., to be published)) is present at the interface of the two protomers at the central axis of the enzyme (Figure 2T). Residues in the N-finger (R4, K5, A7) and β9 strand (V125, Y126, Q127) from both protomers surround the ligand within the binding site. The side chains of R4 and from one protomer and Q127 from the second protomer make hydrogen bonds with the urea nitrogen atoms.

```
SARS_CoV2     1    SGFRKMAFPSGKVEGCMVQVTCGTTTLNGLWLDDVVYCPR
SARS_CoV      1    SGFRKMAFPSGKVEGCMVQVTCGTTTLNGLWLDDTVYCPR
MERS_CoV      1    SGLVKMSHPSGDVEACMVQVTCGSMTLNGLWLDNTVWCPR

SARS_CoV2     41   HVICTSEDMLNPNYEDLLIRKSNHNFLVQ---AGNVQLRV
SARS_CoV      41   HVICTAEDMLNPNYEDLLIRKSNHSFLVQ---AGNVQLRV
MERS_CoV      41   HVMCPADQLSDPNYDALLISMTNHSFSVQKHIGAPANLRV

SARS_CoV2     78   IGHSMQNCVLKLKVDTANPKTPKYKFVRIQPGQTFSVLAC
SARS_CoV      78   IGHSMQNCLLRLKVDTSNPKTPKYKFVRIQPGQTFSVLAC
MERS_CoV      81   VGHAMQGTLLKLTVDVANPSTPAYTFTTVKPGAAFSVLAC

SARS_CoV2     118  YNGSPSGVYQCAMRPNFTIKGSFLNGSCGSVGFNIDYDCV
SARS_CoV      118  YNGSPSGVYQCAMRPNHTIKGSFLNGSCGSVGFNIDYDCV
MERS_CoV      121  YNGRPTGTFTVVMRPNYTIKGSFLCGSCGSVGYTKEGSVI

SARS_CoV2     158  SFCYMHHMELPTGVHAGTDLEGNFYGPFVDRQTAQAAGTD
SARS_CoV      158  SFCYMHHMELPTGVHAGTDLEGKFYGPFVDRQTAQAAGTD
MERS_CoV      161  NFCYMHQMELANGTHTGSAFDGTMYGAFMDKQVHQVQLTD

SARS_CoV2     198  TTITVNVLAWLYAAVINGDRWFLNRFTTTLNDFNLVAMKY
SARS_CoV      198  TTITLNVLAWLYAAVINGDRWFLNRFTTTLNDFNLVAMKY
MERS_CoV      201  KYCSVNVVAWLYAAILNGCAWFVKPNRTSVVSFNEWALAN

SARS_CoV2     238  NYEPLT-QDHVDILGPLSAQTGIAVLDMCASLKELLQNGM
SARS_CoV      238  NYEPLT-QDHVDILGPLSAQTGIAVLDMCAALKELLQNGM
MERS_CoV      241  QFTEFVGTQSVDM---LAVKTGVAIEQLLYAIQQLY-TGF

SARS_CoV2     277  NGRTILGSALLEDEFTPFDVVRQCSGVTFQ
SARS_CoV      277  NGRTILGSTILEDEFTPFDVVRQCSG----
MERS_CoV      277  QGKQILGSTMLEDEFTPEDVNMQIMGVV--
```

**Figure S1**: Sequence alignment between SARS-CoV2, SARS-CoV and MERS-CoV M$^{pro}$. The alignment is colored according to the conservation of residues.

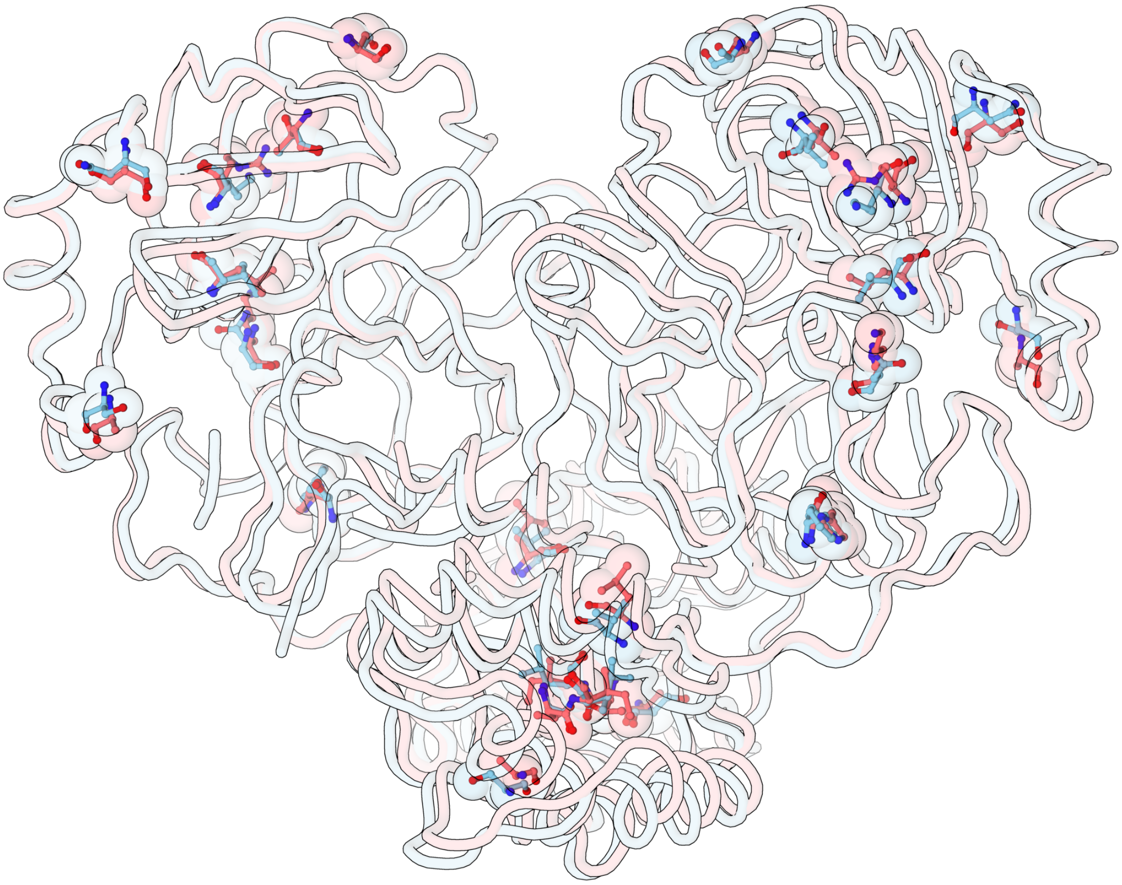

| Residue No. | 35 | 46 | 65 | 86 | 88 | 94 | 134 | 180 | 202 | 267 | 285 | 286 |
|---|---|---|---|---|---|---|---|---|---|---|---|---|
| **SARS-CoV2** | **V** | **S** | **N** | **V** | **K** | **A** | **F** | **N** | **V** | **S** | **A** | **L** |
| **SARS-CoV** | **T** | **A** | **S** | **L** | **R** | **S** | **H** | **K** | **L** | **A** | **T** | **I** |

**Figure S2**: Superimposition of SARS-CoV2 (cyan) and SARS-CoV (red) $M^{pro}$ structures. The spatial position of the dissimilar residues is highlighted. The table lists the dissimilar residues.

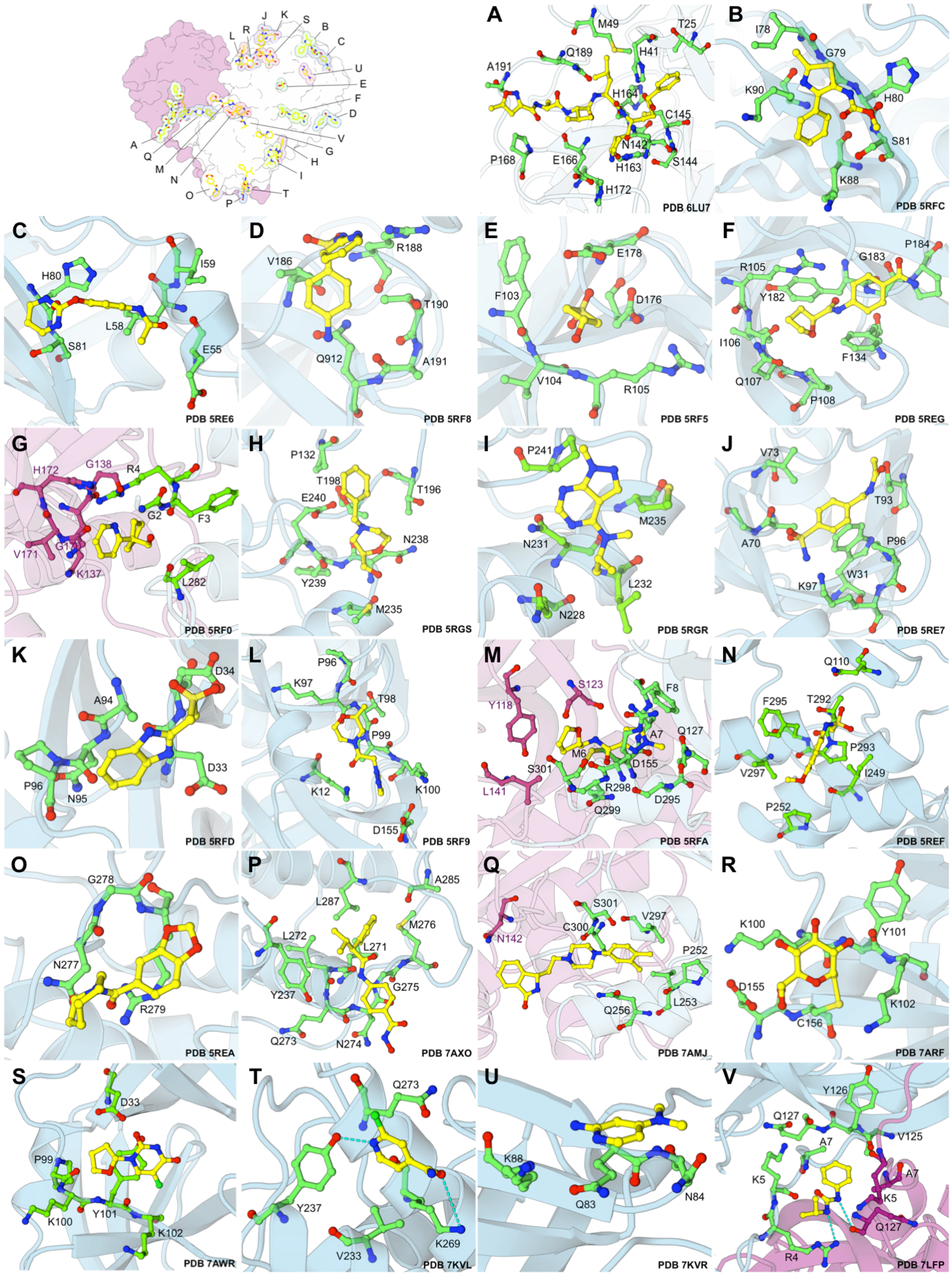


**Figure S3**: Interactions between the SARS-CoV2 (green) and the ligand (yellow) in their representative ligand binding sites A-V. Residues within 4.0 Å of the ligand have been highlighted as green sticks. The protomers are coloured in cyan and pink; the PDB entry of the representative structure is annotated in the bottom right corner. Binding site on only one protomer is illustrated for clarity

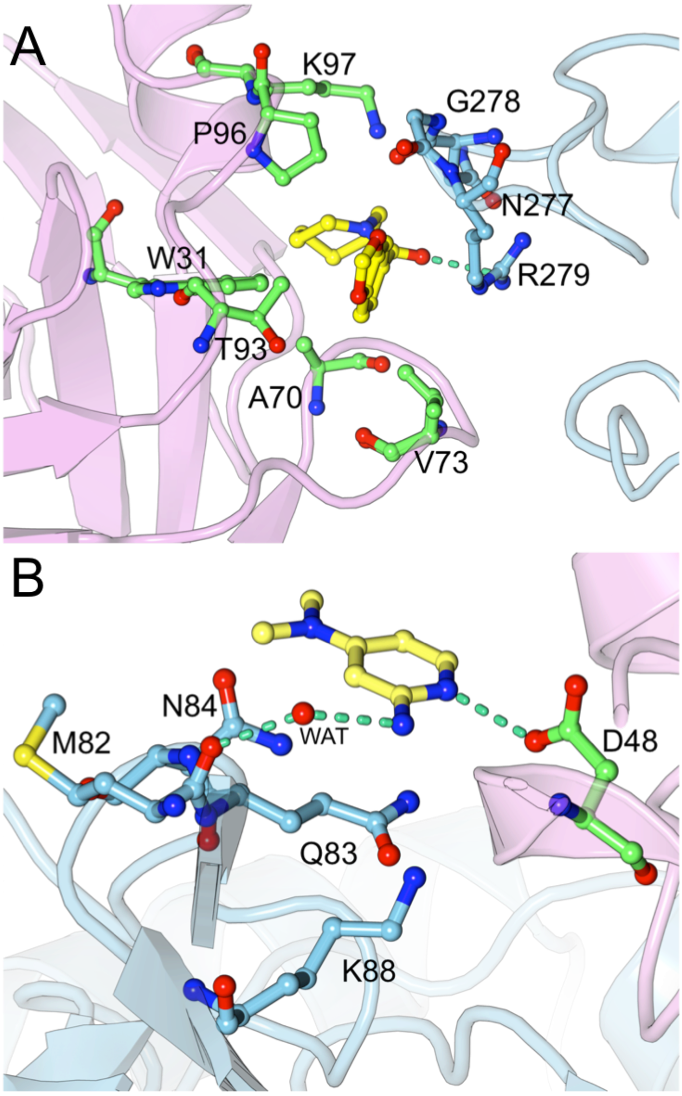


**Figure S4**: Interactions between the SARS-CoV2 M$^{pro}$ (cyan) in the pseudo-ligand binding sites. (A) The ligand (yellow sticks) in site O (cyan sticks) interacts with residues in site J in its symmetry-related protomer (pink ribbon/green sticks) in PDB 5REA. (B) The ligand in site U is stabilized by hydrogen bond interaction with the backbone of M82 via a water molecule and with D48 from its symmetry-related protomer (pink ribbon/green sticks) in PDB 7KVR.

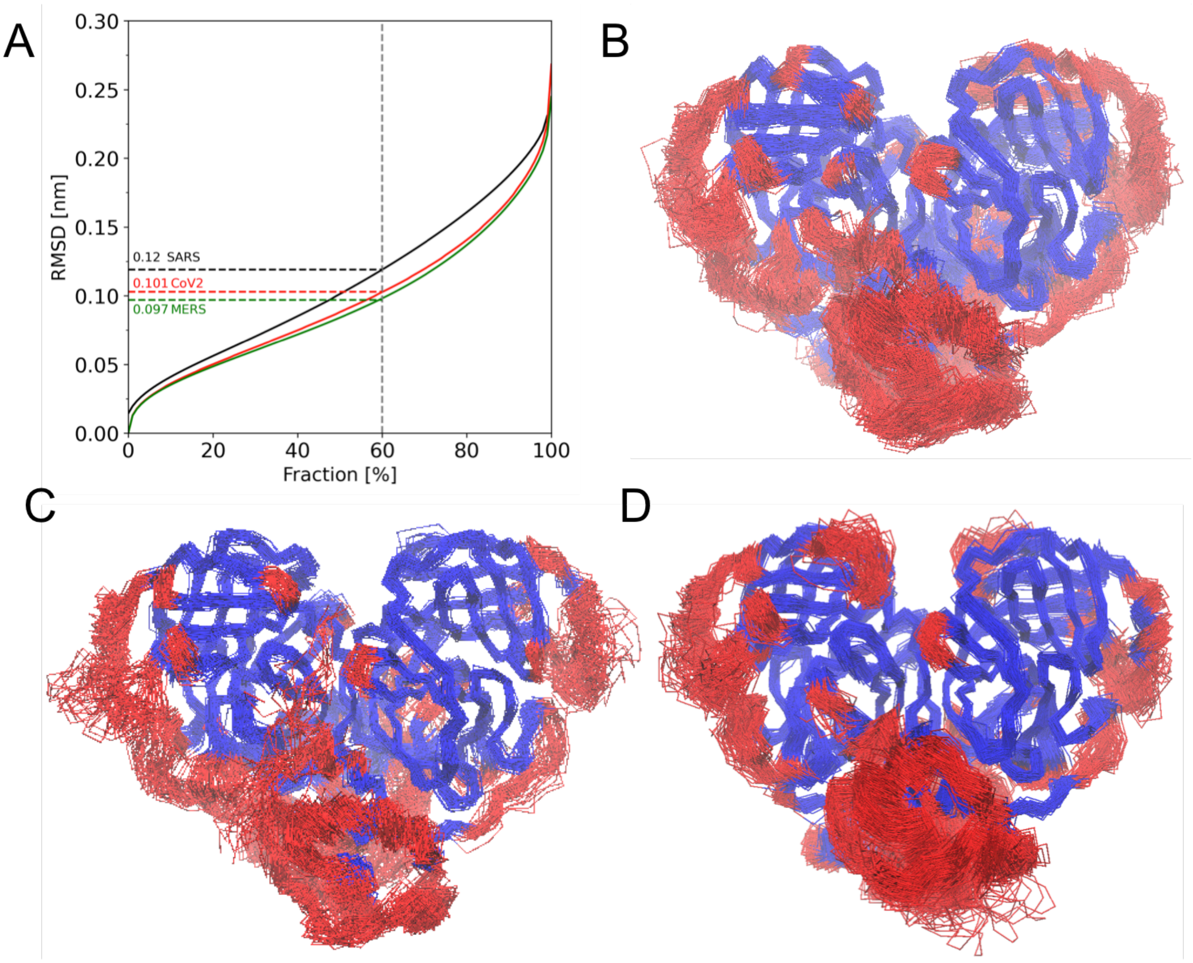


**Figure S5**: Conformational drift in $M^{pro}$ enzymes. (A) 60% of residues in SARS-CoV2 (red), SARS-CoV (black) and MERS-CoV (green), can be aligned to below 0.10 nm, 0.12 nm and 0.09 respectively. Conformations of (B) SARS-CoV2 (C) SARS-CoV and (D) MERS-CoV, extracted at every 100 ns are superimposed to highlight the stable core (blue) and highly dynamic (red) regions in the $M^{pro}$. Domain I and II are most stable, except for the loops and the linker regions. The dimerization domain III is the most dynamic region of the enzyme.

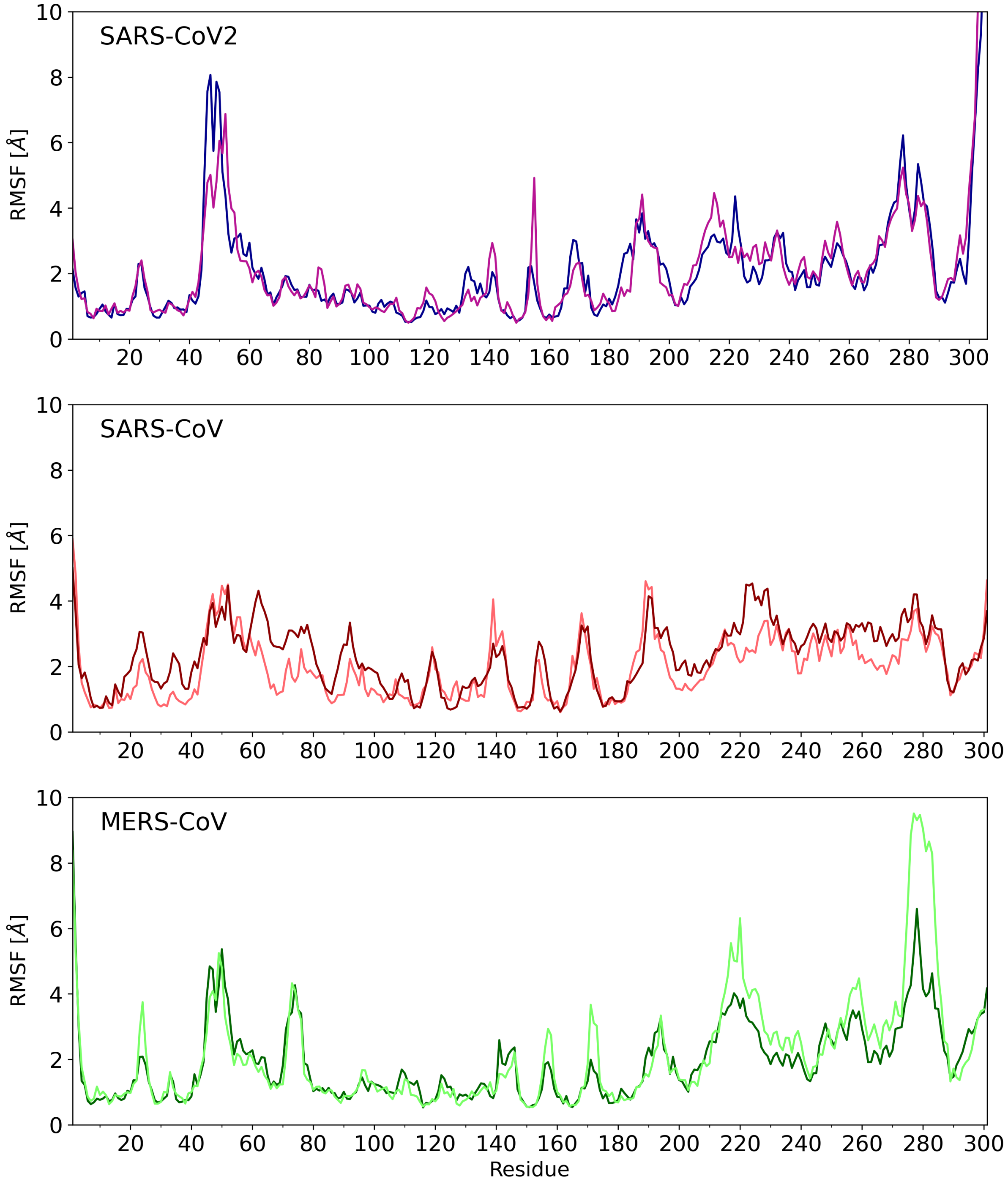


**Figure S6**: Root Mean Squared Fluctuation (RMSF) plots of M$^{pro}$ enzymes. The fluctuations in the both protomers are shown separately.

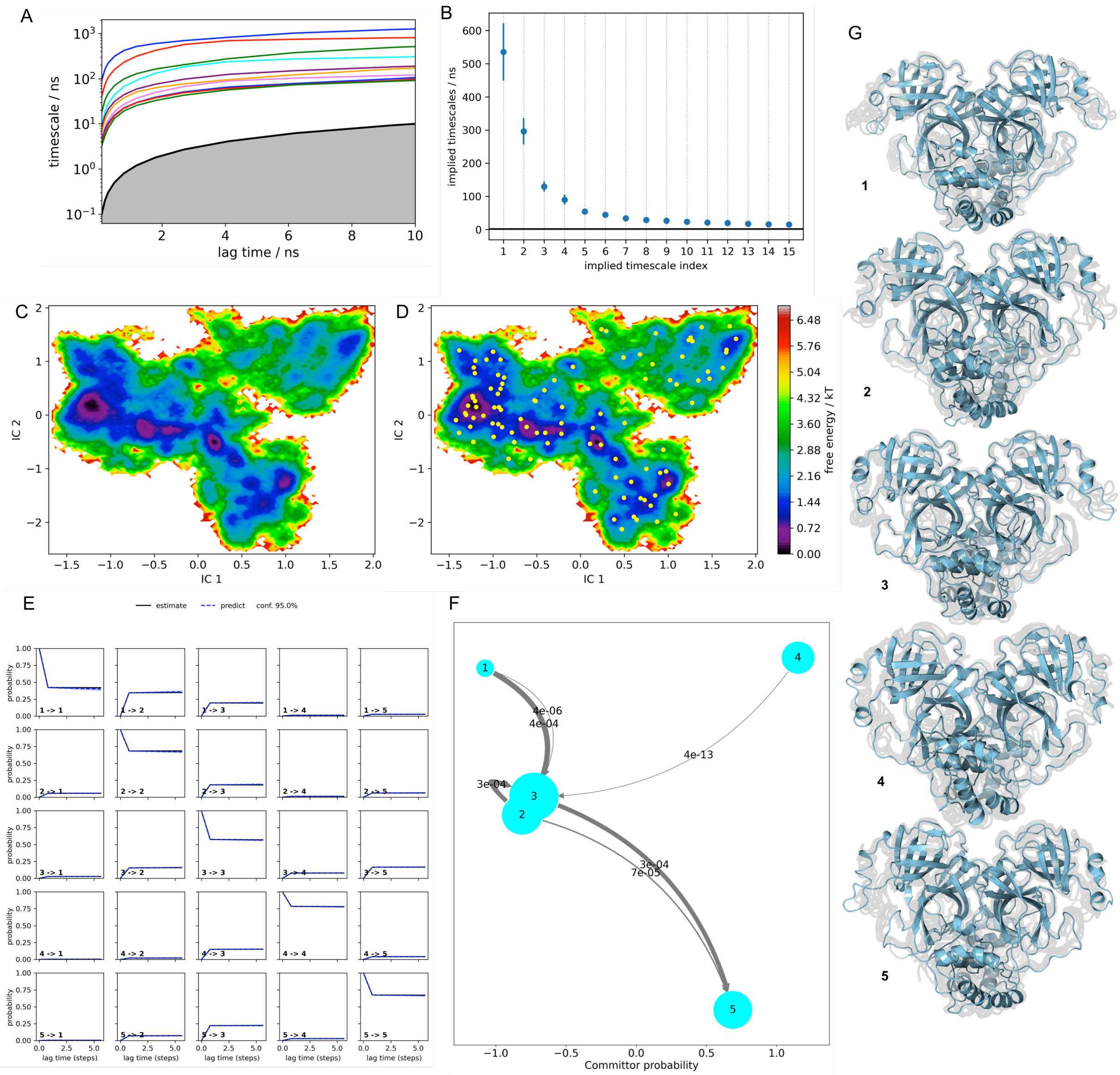


**Figure S7**: Markov State Model of SARS-CoV2 $M^{pro}$ enzyme. (A) Implied timescales (B) Ev/ITS index (C) Free energy landscape (D) Cluster centers plotted on the free energy landscape (E) Chapman-Kolmogorov plot (F) Transition path theory analysis. The state number 1 is the state to which all the starting frames of the trajectories have been aligned. We assume this macrostate to be the crystal structure-encompassing state. The size of the circles are proportional to their population, thus we label it as a dominant state. The fluxes (reactive transition probabilities) are shown as rates (probability per time unit) and shown on the paths. (G) Metastable conformations. The ensemble of backbone geometries contained in each state is illustrated by displaying overlays of the most probably structures of the state (cartoon) on top of samples of the entire state (transparent lines) to highlight both the intrastate conformational variability and the interstate conformational differences. The most probable position of the backbone in each metastable state is illustrated as a cartoon.

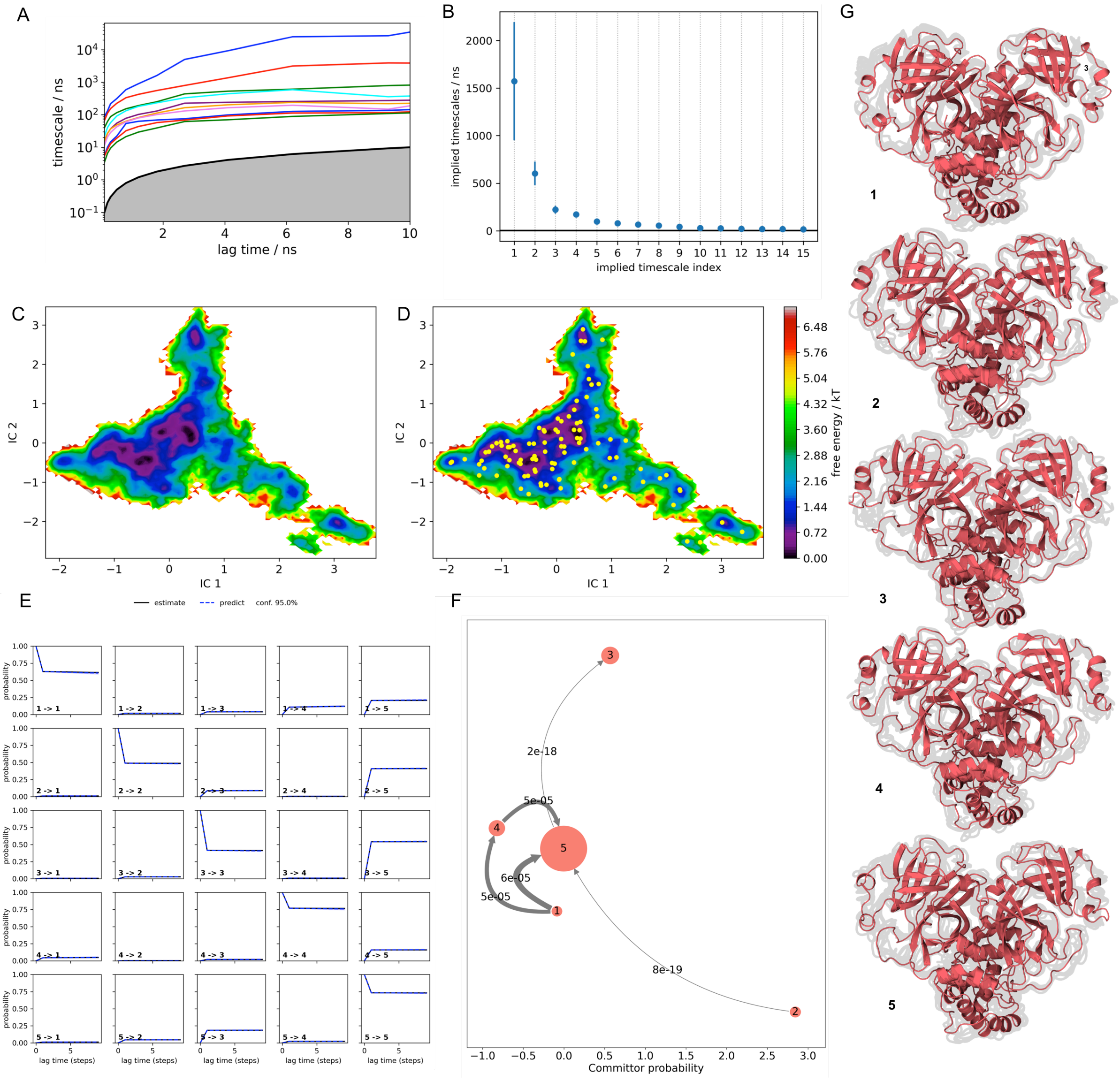


**Figure S8**: Markov State Model of SARS-CoV $M^{pro}$ enzyme. (A) Implied timescales (B) Ev/ITS index (C) Free energy landscape (D) Cluster centers plotted on the free energy landscape (E) Chapman-Kolmogorov plot (F) Transition path theory analysis. The state number 1 is the state to which all the starting frames of the trajectories have been aligned. We assume this macrostate to be the crystal structure-encompassing state. The size of the circles are proportional to their population, thus we label it as a dominant state. The fluxes (reactive transition probabilities) are shown as rates (probability per time unit) and shown on the paths. (G) Metastable conformations. The ensemble of backbone geometries contained in each state is illustrated by displaying overlays of the most probably structures of the state (cartoon) on top of samples of the entire state (transparent lines) to highlight both the intrastate conformational variability and the interstate conformational differences. The most probable position of the backbone in each metastable state is illustrated as a cartoon.

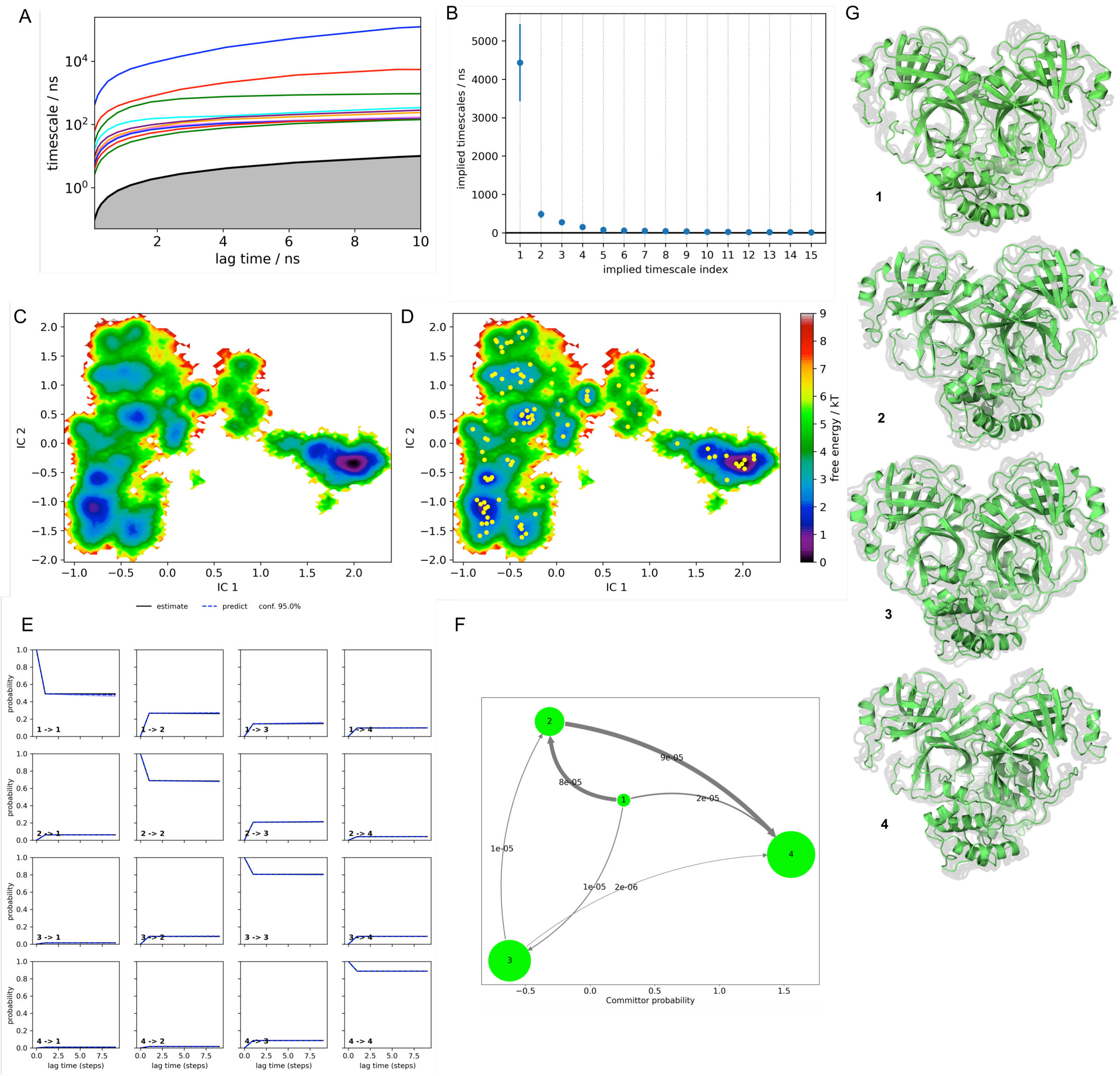


**Figure S9**: Markov State Model of MERS-CoV M$^{pro}$ enzyme. (A) Implied timescales (B) Ev/ITS index (C) Free energy landscape (D) Cluster centers plotted on the free energy landscape (E) Chapman-Kolmogorov plot (F) Transition path theory analysis. The state number 1 is the state to which all the starting frames of the trajectories have been aligned. We assume this macrostate to be the crystal structure-encompassing state. The size of the circles are proportional to their population, thus we label it as a dominant state. The fluxes (reactive transition probabilities) are shown as rates (probability per time unit) and shown on the paths. (G) Metastable conformations. The ensemble of backbone geometries contained in each state is illustrated by displaying overlays of the most probably structures of the state (cartoon) on top of samples of the entire state (transparent lines) to highlight both the intrastate conformational variability and the interstate conformational differences. The most probable position of the backbone in each metastable state is illustrated as a cartoon.

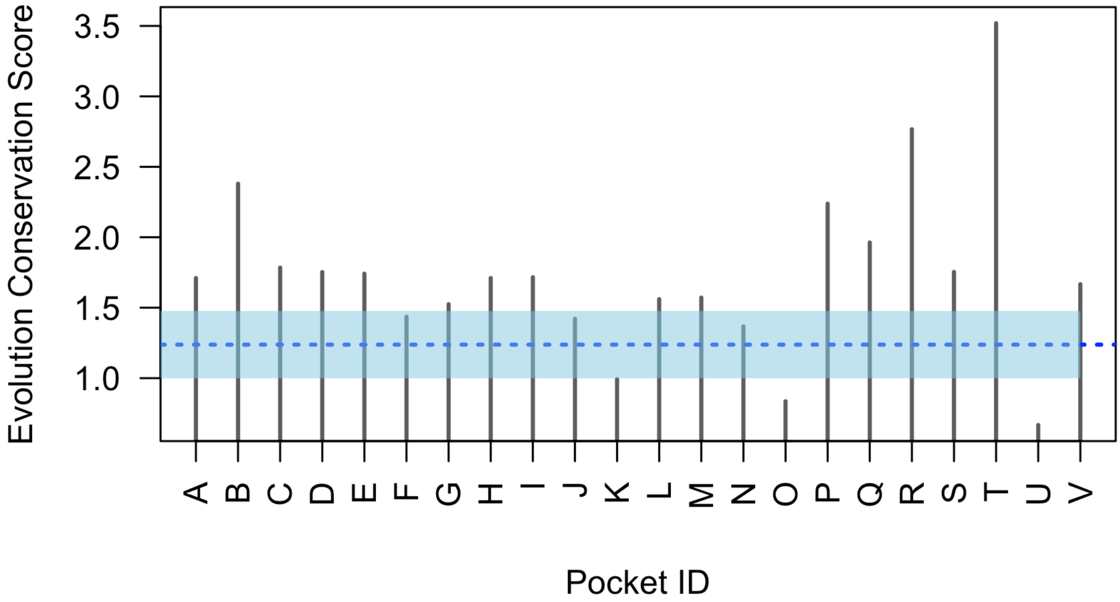


**Figure S10**: Evolution conservation score calculated for each pocket described in Table 1. The score estimates the evolutionary constraints on the pocket as per-residue average of the pairwise correlations. Pairwise correlations are calculated only among pocket residues in the viral MSA (see Methods in the main text). The blue dotted line is the median conservation score calculated from all the surface residue pairs and the blue shaded region highlights the upper and lower limits of the standard deviation.

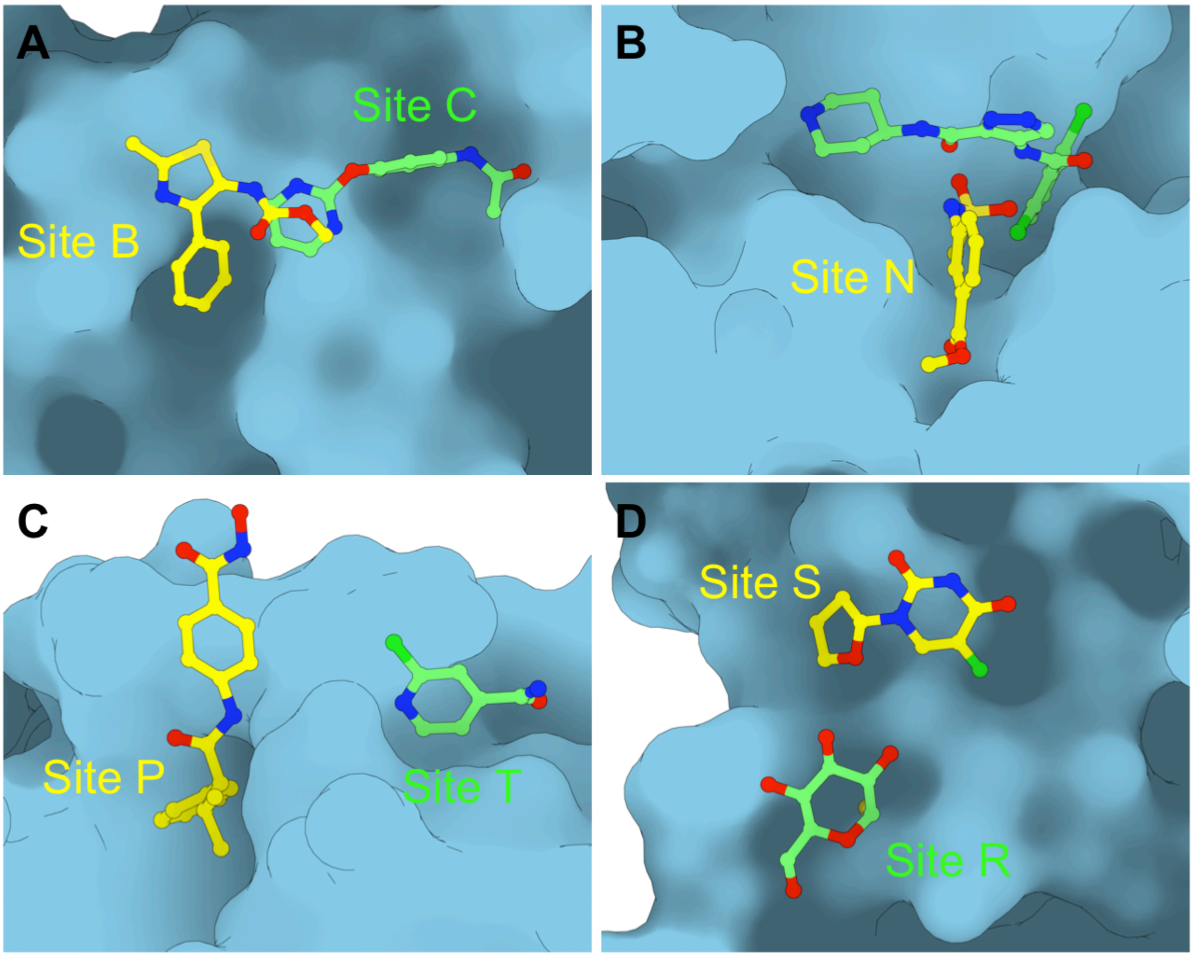


**Figure S11**: Adjacent binding sites in SARS-CoV2.

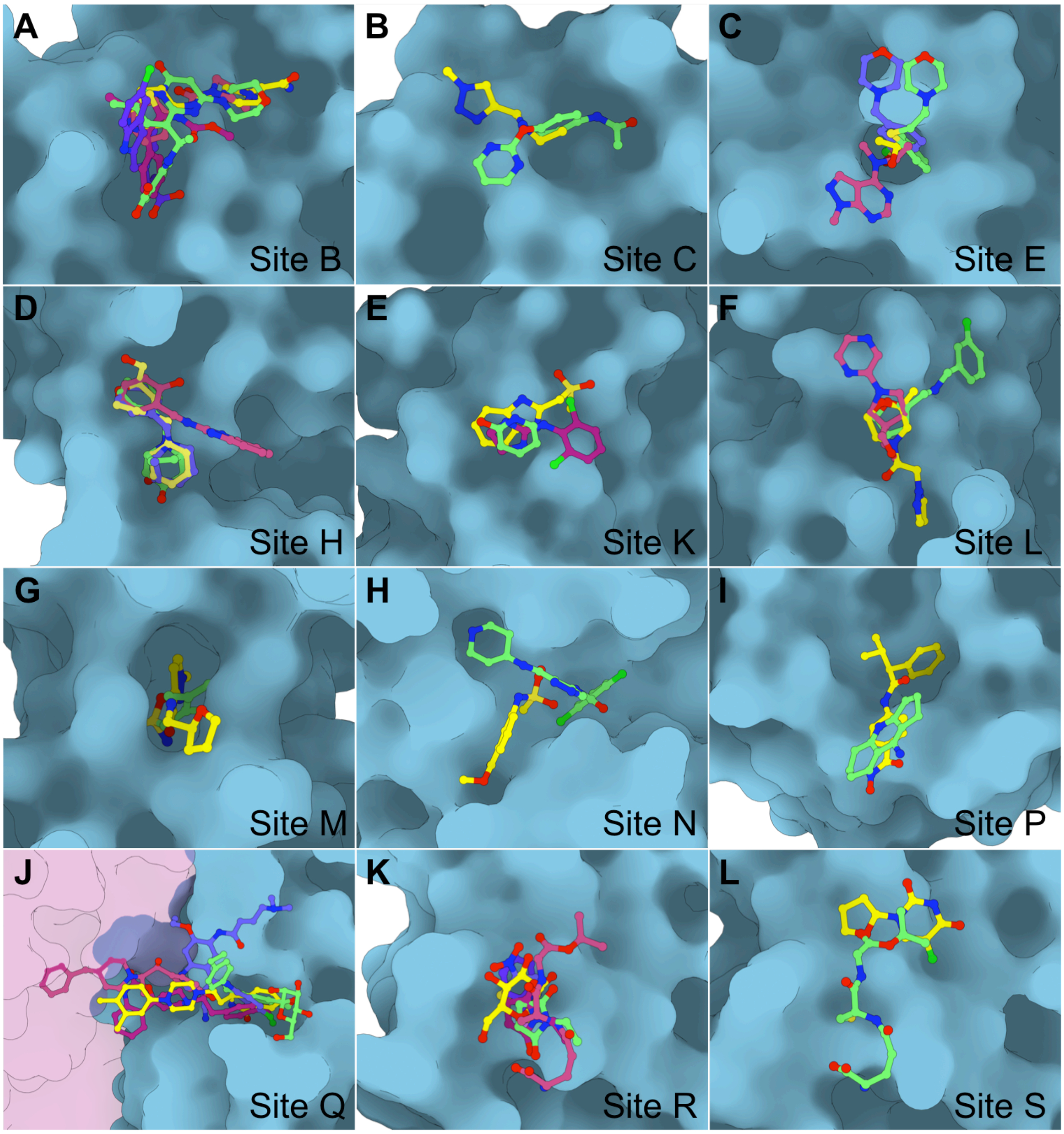


**Figure S12**: Multiple ligands have been identified that bind to (A) Site B, (B) Site C, (C) Site E, (D) Site H, (E) Site K, (F) Site L, (G) Site M, (H) Site N, (I) Site P, (J) Site Q, (K) Site R and (L) Site S. Details of the ligands and their representative PDB are tabled in Table S2. Site A has not been illustrated for clarity.
